# Positive supercoiling buildup is a trigger of *E. coli*’s short-term response to cold shock

**DOI:** 10.1101/2021.12.22.473827

**Authors:** Suchintak Dash, Cristina S.D. Palma, Ines S.C. Baptista, Mohamed N.M. Bahrudeen, Bilena L.B. Almeida, Vatsala Chauhan, Rahul Jagadeesan, Andre S. Ribeiro

## Abstract

Adaptation to cold shock (CS) is a key survival skill of gut bacteria of warm-blooded animals. In *E. coli*, this skill emerges from a complex transcriptional program of multiple, timely-ordered shifts in gene expression. We identified short-term, cold shock repressed (CSR) genes by RNA-seq and provide evidence that their variability in evolutionary fitness is low and that their responsiveness to cold emanates from intrinsic features. Given that their single-cell variability in protein numbers increases after CS, we hypothesized that the responsiveness of a large portion of CSR genes is triggered by the high propensity for transcription locking due to positive supercoiling buildup (PSB). We then proposed a model of this phenomenon and, in support, show that nearly half of CSR genes are highly responsive to Gyrase inhibition. Also, their response strengths to CS and Gyrase inhibition correlate and most CSR genes increase their single-cell variability in protein numbers. Further, during CS, the cells’ nucleoid density increases (in agreement with increased numbers of positive supercoils), their energy levels become depleted (while the resolving of positive supercoils is ATP dependent), and the colocalization of Gyrases and the nucleoid increases (in agreement with increased time length for resolving supercoils). We conclude that high sensitivity to PSB is at the core of the short-term, cold shock responsive transcriptional program of *E. coli* and propose that this gene feature may be useful for providing temperature sensitivity to chromosome-integrated synthetic circuits.

## 1. Introduction

*E. coli* is widely found in the gut of warm-blooded animals in all natural habitats. It usually propagates to new hosts when the original host excretes (or perishes) [Phadtare et al., 1999]. For this, it becomes airborne until encountering new hosts. Thus, it will face (sometimes extreme) temperature downshifts. To cope with these, it has evolved a complex transcriptional program involving many genes [Jones et al., 1987; Phadtare et al., 2004]. Their responses are likely subject to regulatory mechanisms yet to be decoded, which are responsible for the implementation of physiological changes that enhance the chances of survival.

As other prokaryotes, *E. coli* halts cell division and undergoes an “acclimation phase”, during which changes occur at a multi-scale level, from heterogeneous changes in the kinetics of transcription [Oliveira et al., 2016; Charlebois et al., 2018] and translation [Giuliodori et al., 2004; Farewell and Neidhardt, 1998; Phadtare et al., 1999; Keto-Timonen 2016; Madrid et al., 2002], up to decreasing in membrane fluidity [Mansilla et al., 2004; Yamanaka 1999] and increasing cytoplasmic viscosity [Oliveira et al 2016; Parry et al, 2014].

Measurements of transcriptomes at non-optimal temperatures revealed broad responses by specific gene cohorts [Phadtare et al., 2004; Arsène et al., 2000]. During cold shock (CS), a small gene cohort has a fast, transient response, another has a long-term response, while most other genes (including essential genes) remain stable [Phadtare et al., 2004]. This diversity of single-gene responses may be explained by the likely existence of multiple causes for their alterations in expression rates during CS. For example, studies using synthetic gene constructs suggest that temperature can affect the kinetics of rate-limiting steps in transcription initiation, such as the closed and open complex formations [Oliveira et al., 2016], and such effects can differ between promoters [Oliveira et al., 2019]. Other studies showed that temperature affects chromosomal DNA compaction [Goldstein et al., 1984, López-García et al., 2000], which is associated with supercoiling buildup [Stuger et al., 2002; Holmes et al., 2000]. Changing supercoiling buildup levels can cause genome-wide disturbances in gene expression [Travers et al., 2005; Dorman, 2006; Dorman et al., 2016; Peter et al., 2004]. Other influences may be indirect, e.g., temperature affects energy-dependent events, such as interactions between nucleoid-associated proteins (NAPs) and chromosomal DNA [Amit et al., 2003] which affect DNA topology, and thus transcription kinetics [Pruss et al., 1989; Liu et al., 1987; Ma et al., 2013].

Changes in DNA supercoiling may be a quick, efficient means to tune gene expression during stresses, including osmotic shifts [Cheung et al., 2003], oxidative stress [Weinstein-Fischer et al., 2000] and starvation [Drlica et al., 1992]. Many promoters of stress-inducible genes (such as virulence genes in pathogenic bacteria) are sensitive to changes in DNA supercoiling [Dorman, 1995; Dorman 1996]. Thus, it is possible that temperature-dependent changes in DNA superhelical density may be responsible for the responsiveness of some cold shock repressed (CSR) genes.

In agreement, a recent study [Oliveira et al, 2019] tracked RNA production at the molecular level by synthetic variants of the Lac promoter. It was shown that, at low temperatures, RNA production kinetics is weaker and noisier when the gene is chromosome integrated than when it is plasmid borne (in plasmids, supercoiling buildup should be much slower due to the annihilation of positive and negative supercoils [Liu et al., 1987]). They also showed the same phenomenon under Gyrase or Topoisomerase I repression, as well as in energy-depleted cells. Finally, by integrating data from [Phadtare et al., 2004] and [Peter et al., 2004] they hypothesized that CSR genes may exhibit atypical supercoiling-sensitivity.

Here, we subjected *E. coli* cells to CS and identified CSR genes by RNA-seq. We then investigated their common features and their transcription factor (TF) regulation during CS (Figure 1, step 1). Next, we used a YFP fusion library [Taniguchi et al., 2010] to measure CS effects on the single-cell protein levels of 30 CSR genes (Figure 1, step 2). Afterwards, we performed RNA-seq, under Gyrase inhibition, to detect positive supercoiling sensitive (PSS) genes and then identify which genes are both CSR and PSS (Figure 1, step 3). We further collected biophysical data on the chromosome structure and cell energy levels, to support the hypothesis that high supercoiling sensitivity (SS) may provide genes with enhanced short-term CS responsiveness (Figure 1, step 4). Finally, we proposed an analytical model of the dynamics of short-term CSR genes with high PSS (Figure 1, step 5).

**Figure 1:**
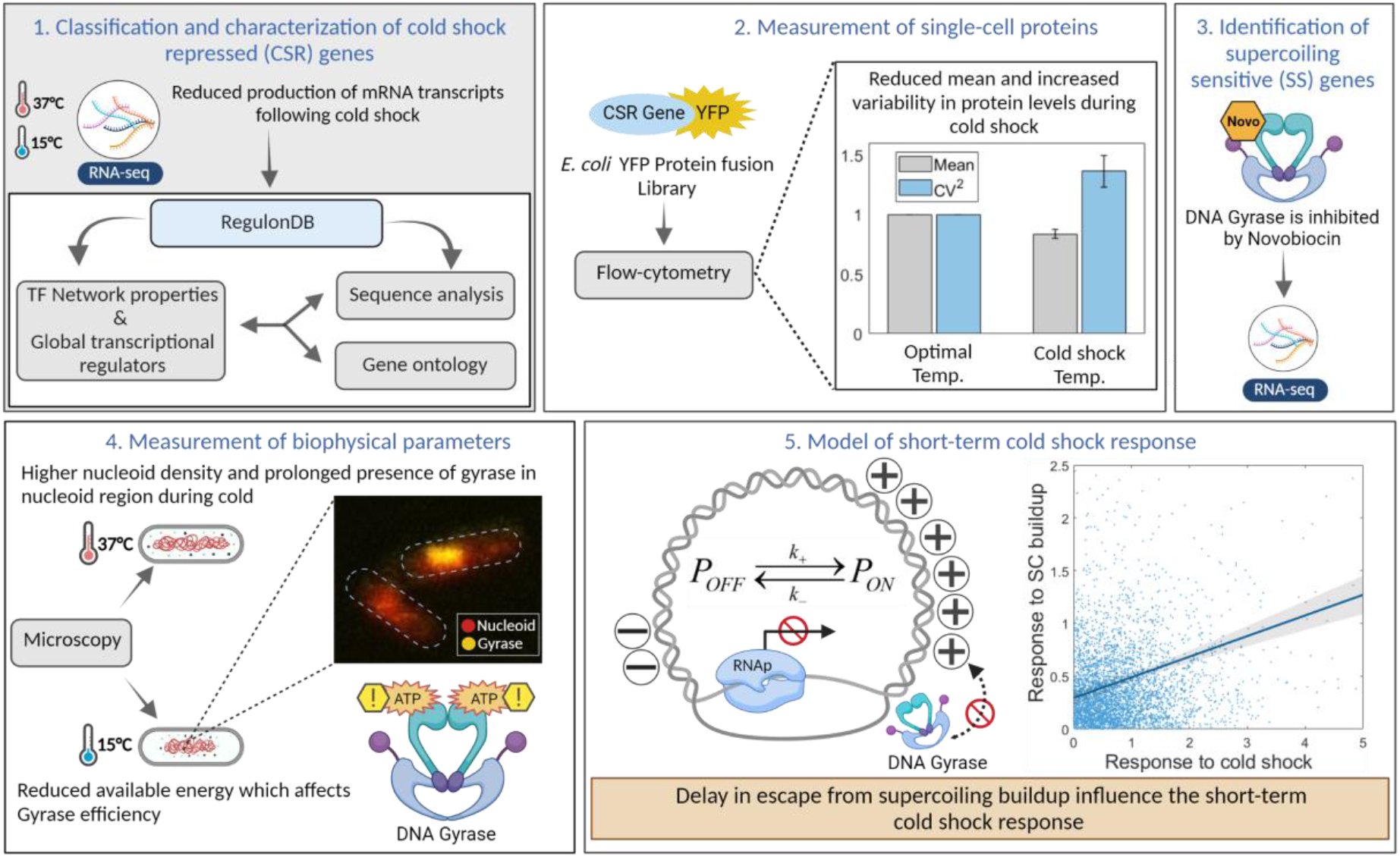
Workflow illustration. **(1)** Identification of short-term CSR genes from RNA-seq data and analysis of their functionality, sequences and, regulation by their direct input TFs and by global transcription regulators (e.g., RNAP). **(2)** Single-cell protein measurements of 30 CSR genes using a YFP fusion library in optimal and CS conditions. **(3)** Identification of PSS genes by RNA-seq following Gyrase inhibition. **(4)** Spectrophotometry and microscopy measurements of biophysical parameters such as ATP levels, cell size, and overlap between the nucleoid and Gyrase. **(5)** Analytical model of short-term CSR due to CS-enhanced locking of promoters due to PSB.

## 2. Material and Methods

### 2.1 Bacterial strains, growth conditions, and gene expression measurements

We used *E. coli* K-12 MG1655 for RNA and protein measurements, since it is the control strain of the YFP fusion library (Supplementary Table S1) [Taniguchi, et al. 2010]. From a glycerol stock (at - 80°C), cells were streaked on LB agar plates and incubated at 37°C overnight. The next day, a single colony was picked from the plate, inoculated in fresh LB medium supplemented with antibiotics (34 μg/mL chloramphenicol for YFP tagged strains) and incubated at 30°C overnight with shaking at 250 RPM. Overnight culture cells were then diluted into fresh M9 media, supplemented with 0.4 % glucose, amino acids, and vitamin solutions, until reaching 0.03 OD (Optical Density at 600 nm measured by Ultrospec 10, Amersham biosciences, UK) and allowed to grow at 30°C with aeration until reaching the mid-exponential phase of growth (OD of 0.3). At this moment, the temperature was downshifted (Innova® 40 incubator, New Brunswick Scientific, USA) and cells were incubated for another 180 mins. Cold-shock conditions are imposed by placing cells at 10-15°C [Phadtare et al., 2004]. Culture temperatures were monitored using a thermometer.

For measurements under Gyrase inhibition (Figure 6D), the antibiotic Novobiocin was added (50 μg/mL) upon reaching the OD ∼0.3. This did not disturb the growth rate (Supplementary Figure S6). To measure RpoS, we used a MG*mCherry* (*rpoS*:mCherry) strain (kind gift from James Locke [Patange et al., 2018]), where the *rpoS* gene codes for σ^38^, which is endogenously tagged with mCherry. For intracellular ATP measurements, we used the QUEEN 2m, a kind gift from Hiromi Imamura [Yaginuma et al., 2014] (Supplementary Table S1 for details).

Since all strains used contain the gene acrA, 50 μg/mL of Novobiocin is not expected to affect cell division rate [Ma et al., 1995]. We further verified this by measuring growth rates by OD_600_. In agreement, growth rates only increase for 200 μg/ml or higher (Supplementary Figure S6).

We measured RNA and protein expression levels by RNA-seq (Supplementary Section I) and by Flow-cytometry (Supplementary Section II), respectively. We used pulse width data from flow-cytometry as a proxy for cell volume [Bahrudeen MNM et al., 2019; Cunningham et al., 1990; Traganos et al., 1984], which assisted the estimation of protein concentrations. We verified these results using microscopy data and image analysis (Supplementary Section III).

Finally, we measured the regions occupied by Gyrase and RNAP, using strains from the YFP fusion library, as described in Supplementary Section III.

### 2.2. Nucleoid visualization by DAPI

To study the effect of cold shock on nucleoid size (Figure 7A), cells were fixed with 3.7% formaldehyde in phosphate-buffered saline (PBS, pH 7.4) for 30 min at room temperature, followed by washing with PBS to remove excess formaldehyde. The pellets were suspended in PBS, and DAPI (4′,6-diamidino-2-phenylindole) (2 μg/mL) was added to the suspension to stain the nucleoid. After incubating for 20 min in the dark, cells were centrifuged and washed twice with PBS to remove excess DAPI. Cells were then re-suspended in PBS and 3 μL of these cells were placed on a 1% agarose gel pad for microscopy [Chazotte et al., 2011].

### 2.3. Cellular ATP levels

QUEEN-2m cells (Supplementary Table S1) were grown as described in Methods Section 2.1. We tracked ATP levels (Supplementary Figure S11) using a Biotek Synergy HTX Multi-Mode Reader (spectrophotometer). The solution was excited at 400 nm and emission was recorded at 513 nm. Similarly, the solution was re-excited at 494 nm and emission was recorded at 513 nm. The ratio of 513 nm emission intensity at these 2 excitation wavelengths, denoted as “400ex/494ex”, is used to quantify cellular ATP concentration as proposed in [Yaginuma H et al., 2014].

### 2.4. Stochastic Model of cold shock response

We used stochastic simulations to estimate the expected noise in gene expression (as measured by the squared coefficient of variation, CV^2^, of gene expression levels in individual cells), assuming the models described in Figure 6A. Simulations were performed using SGNSim [Ribeiro et al., 2007], whose dynamics follows the Stochastic Simulation Algorithm [Gillespie, 1976; Gillespie, 1977]. The time length of each simulation was 10^6^s, which sufficed to avoid fluctuations due to sources other than noise in gene expression [Häkkinen et al., 2016]. The results [ Figure 6B] were collected from 100 runs for each model, which sufficed to obtain consistent results. Finally, at the start of each run, in addition to the parameter values in Supplementary Table S8, it was set that there is 1 promoter in the system. In Model 1.3 (Figure 6A), the promoter was initially in the “ON” state.

### 2.5. Information from RegulonDB

Our data was extracted from RegulonDB v10.9. The data includes information on TF interactions, operon organization, and nucleotide sequence.

## 3. Results

### 3.1 Cell morphology, physiology, and master transcription regulators during cold shock

Having subject cells to CS (Methods Section 2.1), we first studied physiological and morphological effects. Once at 15°C or lower temperatures, cells no longer divided (Figure 2A). Meanwhile, their size was not affected, according to microscopy (Supplementary Figure S7B) and flow-cytometry (Figure 2C, Supplementary Section II) data. Nevertheless, these cells are not likely to be shifting to stationary growth, since RpoS concentrations remain low (Supplementary Figure S15) [Lange, 1991; Jishage et al 1996] (Methods Section 2.1), when compared to cells in optimal conditions and to cells in the stationary growth phase (Figure 2B).

**Figure 2.**
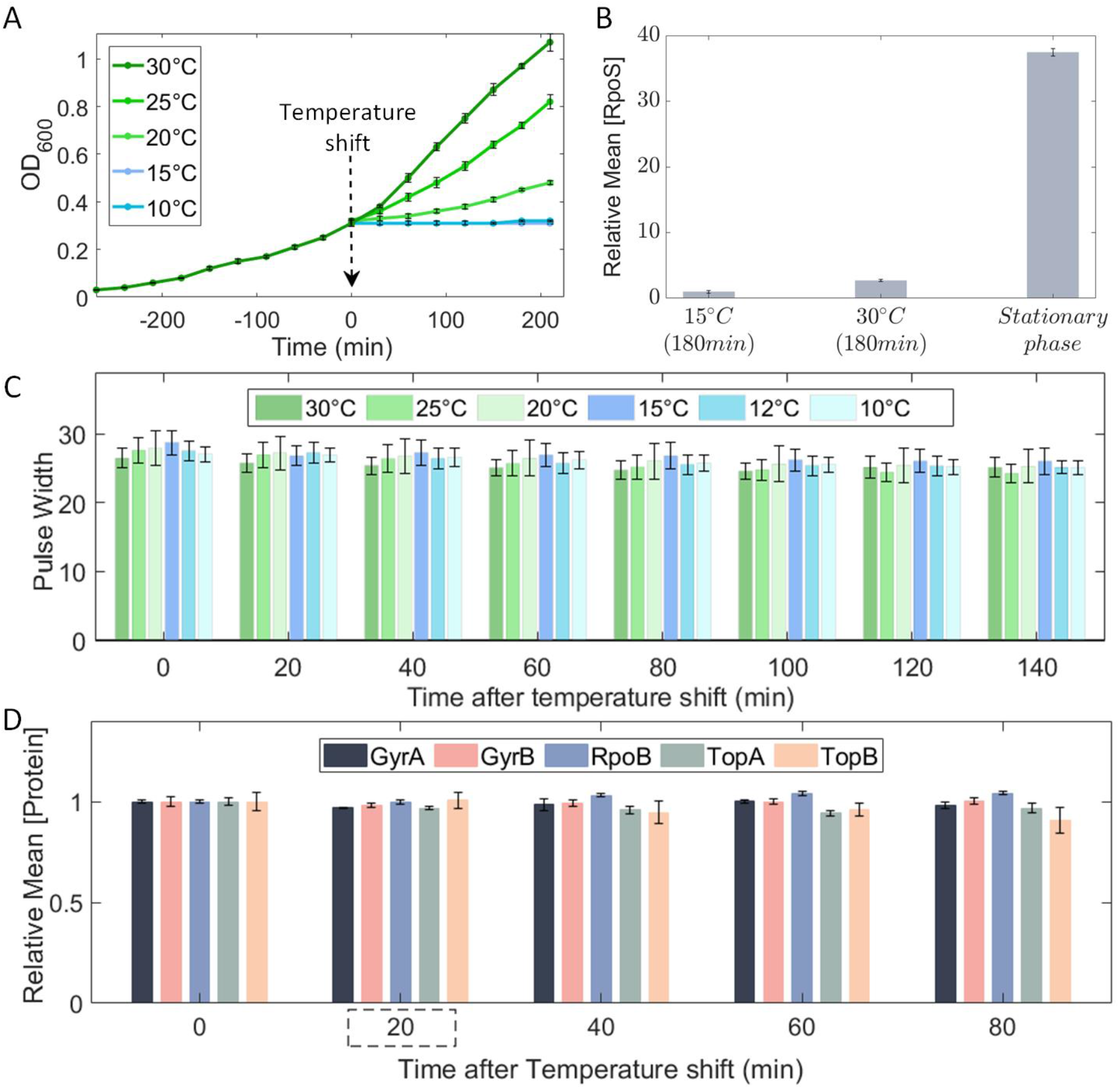
Effects of temperature shifts on cellular morphology, physiology, and global transcriptional regulators. **(A)** Growth curves at 10°C, 15°C, 20°C, 25°C, and 30°C following a temperature shift, set to be minute 0. **(B)** Mean RpoS concentration during CS and optimal conditions after 180 min, and during stationary growth (i.e., after 700 min). **(C)** Pulse width over time following temperature shifts (Methods Section 2.1). **(D)** Mean concentration of GyrA, GyrB, TopA, TopB and RpoB proteins over time after shifting temperature to 15°C. The vertical error bars are the standard error of the mean (SEM) from 3 biological repeats.

Next, we examined potential short-term effects of CS on the concentrations of the master regulators of transcription, since if they change, it could influence the dynamics of CSR genes. In detail, we observed RNA polymerase (RNAp) by tracking a YFP tagged β subunit, which is the product of the rpoB gene (Supplementary Table S1). We also observed the two subunits of Gyrase (GyrA and GyrB) and of Topoisomerase I (TopA and TopB) using a YFP fusion library [Taniguchi et al., 2010], since they are the master regulators of DNA supercoiling levels [Gellert et al., 1976; Wang, et al., 1971]. As such, they heterogeneously influence transcription at a genome-wide level. Further, evidence suggests that the efficiency of Gyrase and Topoisomerase is temperature sensitive [Drlica 1992; Wang et al., 1985; Oliveira et al., 2019].

Neither of these global regulators showed concentration changes during 80 mins after CS (Figure 2D), while the RNA-seq measurements reported below to identify CSR genes were performed 20 min after CS. As such, short-term CS responsiveness, is not expected to be activated by changes in the concentrations of these master transcription regulators.

### 3.2 Identification of short-term CSR genes

We performed RNA-seq measurements (Supplementary Section I) at 0, 20, 80, and 180 min after shifting temperature to 15°C and under optimal (control) temperature (Methods Section 2.1).

We classified single-gene responses to CS as ‘short-term’ when they occur *prior to* influence from direct input TFs or global regulators, or from cell division. As such, based on cell doubling times (Figure 2A) and on known rates of transcription and translation in *E. coli* (see e.g. [Bernstein et al., 2002; Taniguchi et al., 2010]) we expect the changes in RNA numbers at 20 min. after the CS to be short-term, while subsequent changes at 80 and 180 min are here classified as being mid-term and long-term changes, respectively.

To identify short-term CSR genes, we obtained the RNA log_2_ fold changes (LFC_CS_) at 20 mins after shifting to cold shock. We also obtained control LFCs (LFC_CTRL_) after the same time interval when not shifting temperature.

We classified a gene as ‘CSR’ when its LFC_CS_ < 0 (with p-value < 0.05), provided that its corresponding LFC_CTRL_ ≥ 0 (with p-value < 0.05) as this enhances the chance that the repression at CS was due to the CS. We found that 381 genes (Supplementary File X2) respected these conditions and, using the YFP fusion library [Taniguchi et al., 2010], one can measure the proteins levels of 124 of them.

From these 124, we selected genes that: i) have high expression under optimal conditions (resulting in higher fluorescence than cell backgrounds) and; ii) LFC_CS_ < - 0.23, i.e., their RNA levels were reduced by 15% or more, relative to the same RNA in the control condition, to ensure significant downregulation during CS at the protein level. We found 30 of the 124 genes respected these conditions and, thus, we selected them for single-cell fluorescence measurements in the control and CS conditions.

Finally, we selected 6 of these 30 genes and additionally collected single-cell, time-lapse flow-cytometry data on their dynamics. Taken together, their expression levels cover the state space of protein expression levels of the 30 CSR genes.

### 3.3 Ontology and evolutionary fitness of short-term CSR genes

We investigated the ontology [Ashburner et al., 2000; Gene Ontology Consortium, 2021] of CSR genes to identify the most affected biological processes. From an over-representation test (Supplementary Section XVI), we compared the number of CSR genes related to specific biological processes (quantified by the fold enrichment) with the *expected* number, given genome-wide numbers.

The significantly over-represented biological processes are listed in Table S11. Visibly, of 30 major biological processes in *E. coli* considered in GO studies [Ashburner et al. 2000; Gene Ontology Consortium, 2021], CSR genes are mainly associated with metabolism and response to external stimulus (Supplementary Figure S14). This agrees with reports that genes involved in metabolism are commonly affected during CS, which reduces growth rate and the rate of glycolysis [Gadgil et al., 2005; Andersen et al., 1980; Phadtare et al 2004].

Next, we studied the evolutionary fitness of the responsive genes (Supplementary section XVII). Interestingly, while their average fitness is the same as expected by chance, their fitness variability is smaller than in same-sized cohorts of randomly selected genes (Figure 3A). This is not because they are over-represented in two functional groups, since the fitness variability of random cohorts with the same distribution of gene functions (164 genes related to metabolism, 41 genes responsive to stimulus, 36 genes in both groups, and 140 with other functions) also have statistically distinguishable fitness variability from CSR genes. Given that the fitness is positively correlated to the evolutionary conservation (Supplementary Section XVII), we hypothesize that their evolutionary ages are likely to be more similar than expected by chance as is the fitness.

**Figure 3.**
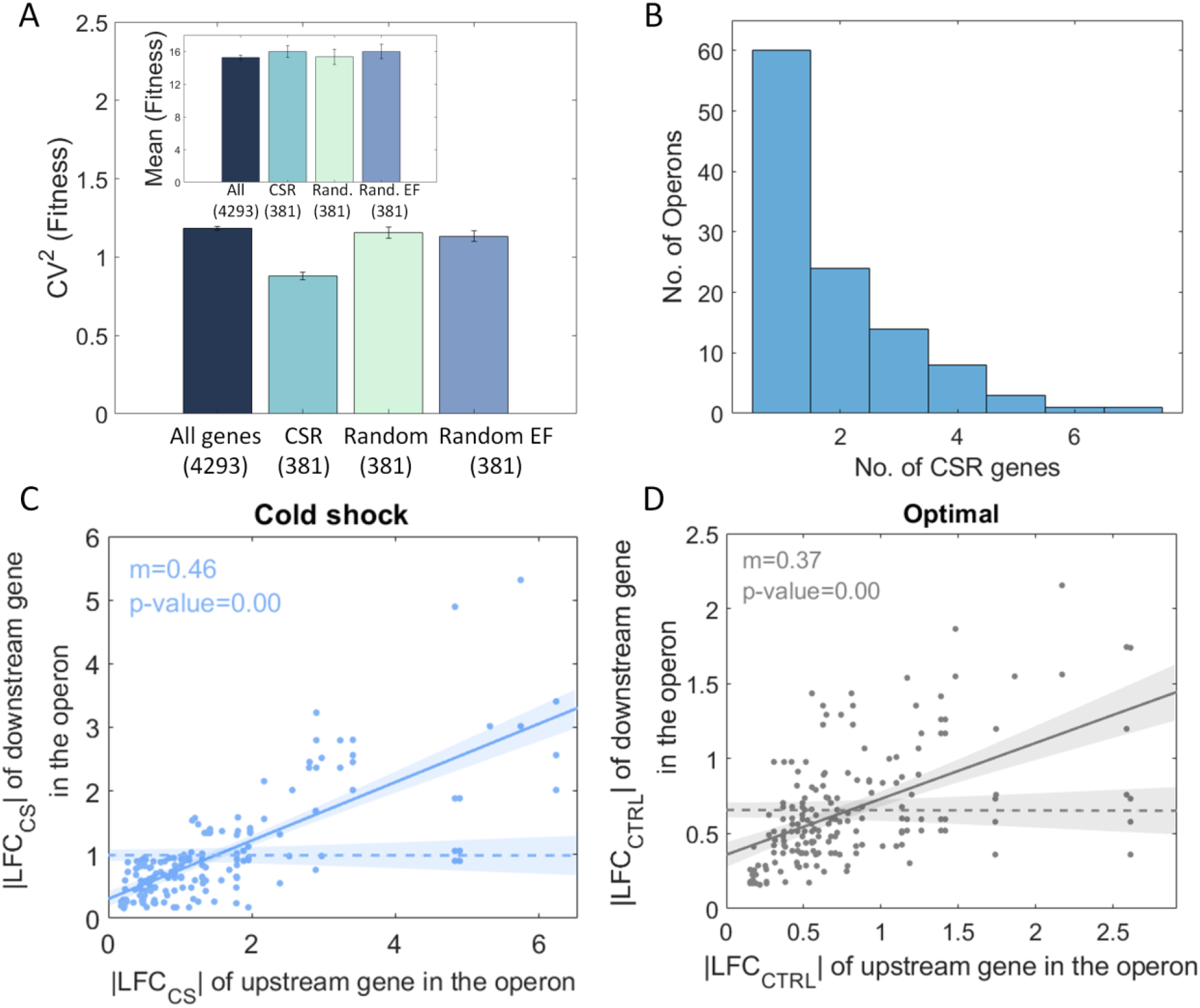
Characterization of CSR genes. **(A)** Bar plot of the variability, CV^2^, of the fitness of all genes of *E. coli*’s genome (dark blue), CSR cohort (light blue), randomly selected cohort (light green) and a randomly selected cohort with the same size and same biological function (purple bar). The inset shows the mean fitness (in %) for each cohort. **(B)** Distribution of CSR genes in operons **(C)** Scatter plot between the | LFC_CS_| of pairs of CSR genes in the same operon during CS. **(D)** |LFC_CTRL_| of CSR genes upstream in the operon plotted against the |LFC_CTRL_| of CSR genes downstream in the same operon at optimal temperatures.

In addition to ontology and fitness, we also investigated the potential influences on CS responsiveness from TFs (Supplementary Section XVIII) and promoters’ AT richness (Supplementary Section XIX). However, we failed to find any relationships.

### 3.4 Short-term responses of CSR genes can be partially explained by operon organization

Genes in the same operon commonly exhibit co-expression [Jacob, F. and Monod, J, 1961; Sabatti et al, 2002]. To verify if this influences the composition of the identified population of CSR genes, we confronted the positionings of CSR genes in the same operons with correlations in their dynamics.

Of the 381 CSR genes, 169 are not in operon structures (according to RegulonDB [Santos-Zavaleta A et al., 2019]), while the remaining 212 are organized in a total of 111 operons (Figure 3B, Supplementary File X3). As expected, the LFC_CSR_ of pairs of the CSR genes in the same operon are (similarly) correlated in both optimal and CS conditions (Figures 3C and 3D).

We confronted these data with a null model that assumes the same distribution of genes per operon as in Figure 3B, but with the genes composing those operons being randomly selected from the set of genes in operons. The random pairs showed no dynamic correlation (Figures 3C and 3D). We conclude that the operons’ organization might partially explain the numbers of CSR genes of *E. coli*, i.e., some genes might be CSR because they are located downstream a CSR gene in the same operon.

Nevertheless, there are 60 operons with only 1 CSR gene (Figure 3B). Thus, for a gene to be CSR, it does not suffice to be in the same operon as a CSR gene.

### 3.5 The scaling between noise and mean of single-cell CSR protein numbers is temperature dependent

Given that the short-term response of CSR genes was uncorrelated with their input TFs dynamics (Supplementary Section XVIII), individual gene features are more likely to be responsible for their repression during cold shock. We expect that, by repressing gene expression at the transcription level, these mechanisms will affect how noise and mean expression relate [Peccoud et al., 1995; Golding et al., 2005; Taniguchi et al., 2010]. To investigate this, we studied the single-cell distributions in protein numbers of 30 CSR genes (Methods Section 2.1, Supplementary Table S1, Supplementary File X1).

To quantify single-cell protein numbers, we first corrected the statistical moments of the distributions to account for cell auto-fluorescence (Supplementary Section IV). Then, we plotted the mean expression levels in optimal conditions against the corresponding protein numbers reported in [Taniguchi et al., 2010] (Supplementary Figure S3). Given the best fitting line, from here onwards we convert protein expression levels into protein numbers using a scaling factor of 0.1. Meanwhile, we did not find correlations between protein levels and cell size (Supplementary Figure S11A), in agreement with the lack of change in cell size with CS (Figure 2C and Supplementary Figure S7) implying that cell size is not affecting single-cell expression levels. Finally, as expected from the mechanical coupling between transcription and translation in *E. coli* (Miller et al., 1970), the changes with CS in these 30 protein numbers correlated to the changes in the corresponding RNA numbers (Supplementary Figure S11B, Supplementary section XI), indicating that protein levels can be used to study the effects of regulatory mechanisms of transcription.

We thus plotted the mean single-cell protein numbers of CSR genes, M, against the corresponding noise, as measured by CV^2^, for each gene and best fitted the data by Ordinary Least Squares (OLS) with the function [Bar-Even et al., 2006; Taniguchi et al., 2010]:

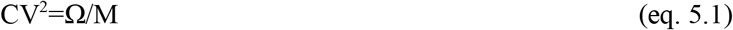

where Ω is a constant and M are mean protein numbers (estimated in Supplementary Figure S3). From Figure 4A, the inverse proportionality between CV^2^ and M, previously observed in optimal conditions [Bar-Even et al., 2006; Taniguchi et al., 2010; Newman et al., 2006], is valid during CS, but CV^2^ becomes higher for the same M (Ω ∼26% higher than in optimal conditions). Meanwhile, since Ω does not change from 120 min to 180 min after the cold shock, changes likely occurred prior to 120 min (Supplementary Figure S2).

**Figure 4.**
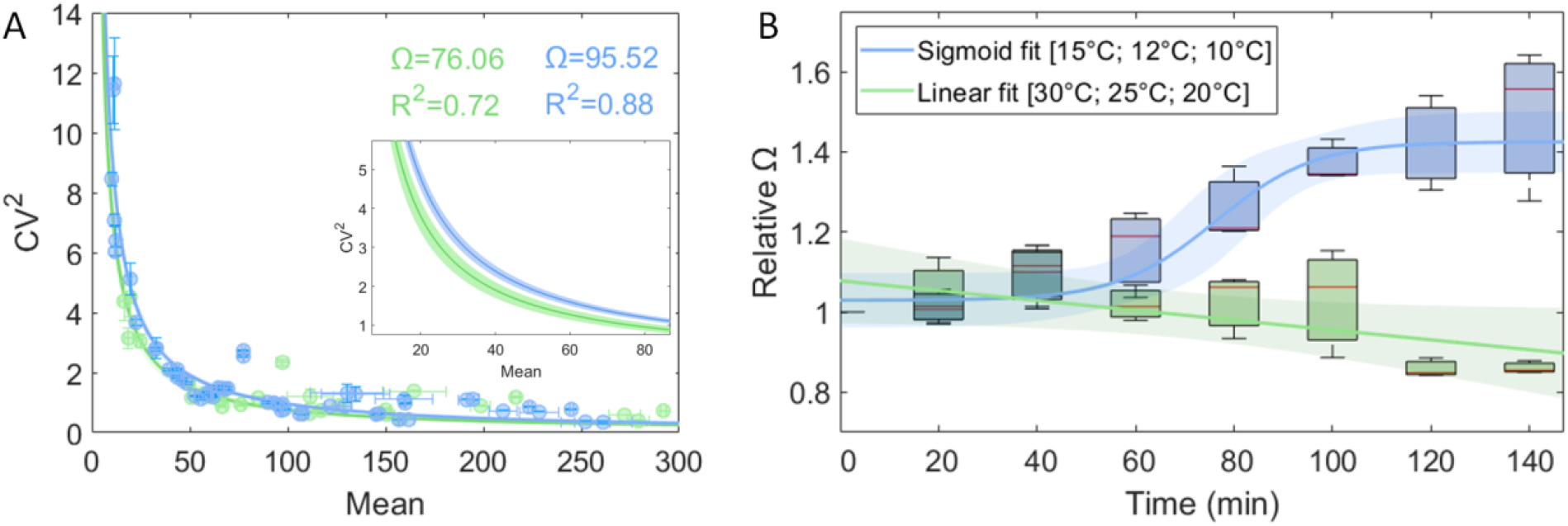
Relationship between CV^2^ and mean protein numbers over time, at different temperatures. Blue corresponds to cold shock conditions, while green corresponds to optimal conditions. **(A)** Squared coefficient of variation (CV^2^) versus mean protein numbers of 30 CSR genes (Supplementary file X1). Data at 120 and 180 min was merged as they did not differ (Supplementary Figure S2). We performed a 2-sample t-test to test the null hypothesis that there is no difference between Ω at 30°C and 15°C. The test rejected the null hypothesis with a p-value of 0.02. **(B)** Box plot of Ω over time at ‘control’ and ‘cold shock’ temperatures. The red line in the box is the median and the top and bottom of the box are one standard deviation (STD) above and below the median, respectively. For control and cold-shock temperatures, we fit the best fitting function. We performed an F-test on the regression model, which tests for the hypothesis that the 0 order polynomial fits significantly better than a 1^st^ order polynomial. The test did not reject the null hypothesis that the best fit line is a horizontal line (p-value of 0.06). The lines correspond to the best-fit functions that maximize R^2^.

To further investigate how Ω changed following CS, we measured single-cell distributions of protein levels of 6 genes each 20 min for 140 min following the temperature shifts. These genes (aldA, feoA, manY, ndk, pepN, tktB) have mean protein levels that cover the state space of M of the 30 CSR genes. For each time moment, we extracted the corresponding Ω that best fits the data (Figure 4B, Supplementary Figure S4). Visibly, Ω increases with time during CS, but not at optimal temperatures (Supplementary Figure S16).

Namely, at T ≤ 15 °C, 40 min after CS, there is a sharp increase in Ω, while at T > 15 °C, Ω remains constant. In detail, for CS temperatures (10 °C, 12 °C and 15 °C), the data is best fit by a sigmoid curve (R^2^= 0.96) of the type 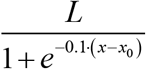, where *L* is the curve’s maximum value and *x*_*0*_ is the value of the sigmoid midpoint (we also attempted to fit polynomials up to the several order, but none fitted better). Meanwhile, for the set of control temperatures, the data is best fit by a first order polynomial (R^2^= 0.91).

Overall, we suggest that, as CS is applied, a step emerges in transcription that is responsible for the strong repression of CSR genes, which not only reduces expression levels of CSR genes, but it also increases the scaling factor between noise and mean of protein numbers.

Finally, from [Taniguchi et al., 2010], most protein number distributions in optimal conditions are well described by a Γ distribution. In these, both (eq. 5.1) is valid, as well the skewness can also be written as a function of M (Derivation in Supplementary Section Va) as follows:

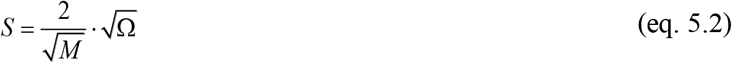

Given the Ω values above, we estimated S using (eq. 5.2) and compared to the empirical values of S in cold shock and control conditions (Figure 5A-B). We find that the two correlate linearly (see also Supplementary Figure S5), above the noise floor, which was estimated using the data in Figure 4A (Supplementary Figure S10, Supplementary Section XIV). This suggests that the effects of CS propagate up to the third moment of the single-cell distribution of protein numbers.

**Figure 5.**
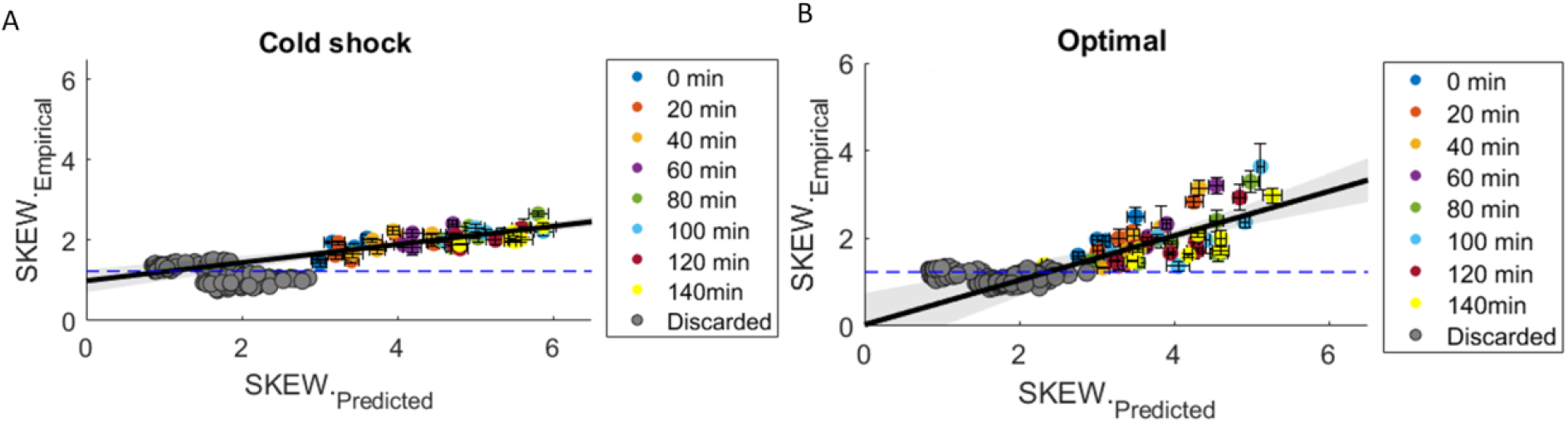
Correlation between empirical and predicted skewness. **(A)** CS temperatures (15°C, 12°C and 10°C). Skewness is predicted using equation 5.2 and the empirical values of Ω (Section 3.5). **(B)** Control temperatures (30°C, 25°C and 20°C). Meanwhile, empirical data on skewness is extracted from single-cell distributions obtained by flow-cytometry (Supplementary File X1) after being corrected for background noise. Blue dashed line is the estimated lower bound (Supplementary Section XIV). Grey circles are data points excluded from the fitting due to being below or crossing the noise floor.

### 3.6 An ON-OFF model can explain the short-term dynamics of CSR genes

From past studies [Taniguchi et al., 2010], *E. coli* transcription in optimal conditions can be well modeled as a one-step process (reaction 1.1 in Figure 6A). Using this reaction, along with reactions for translation (reaction 2 in Figure 6A) and RNA and protein decay due to degradation and dilution in cell division (reactions 3 and 4, respectively), one can model the dynamics of RNA and protein numbers in single-cells. Using this model, we derived an analytical solution for Ω (Supplementary Section Vb):

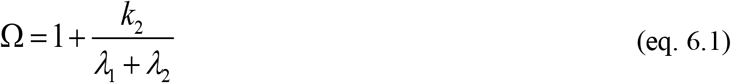

where k_2_ is the translation rate, and λ_1_ and λ_2_ are the RNA and protein decay, respectively. Given this, and since λ_1_ >> λ_2_ [Koch and Levy, 1955; Taniguchi et al., 2010], Ω would then necessarily be controlled by (k_2_ / λ_1_). Because of it, this model cannot explain the selection of the CSR gene cohort during CS.

**Figure 6.**
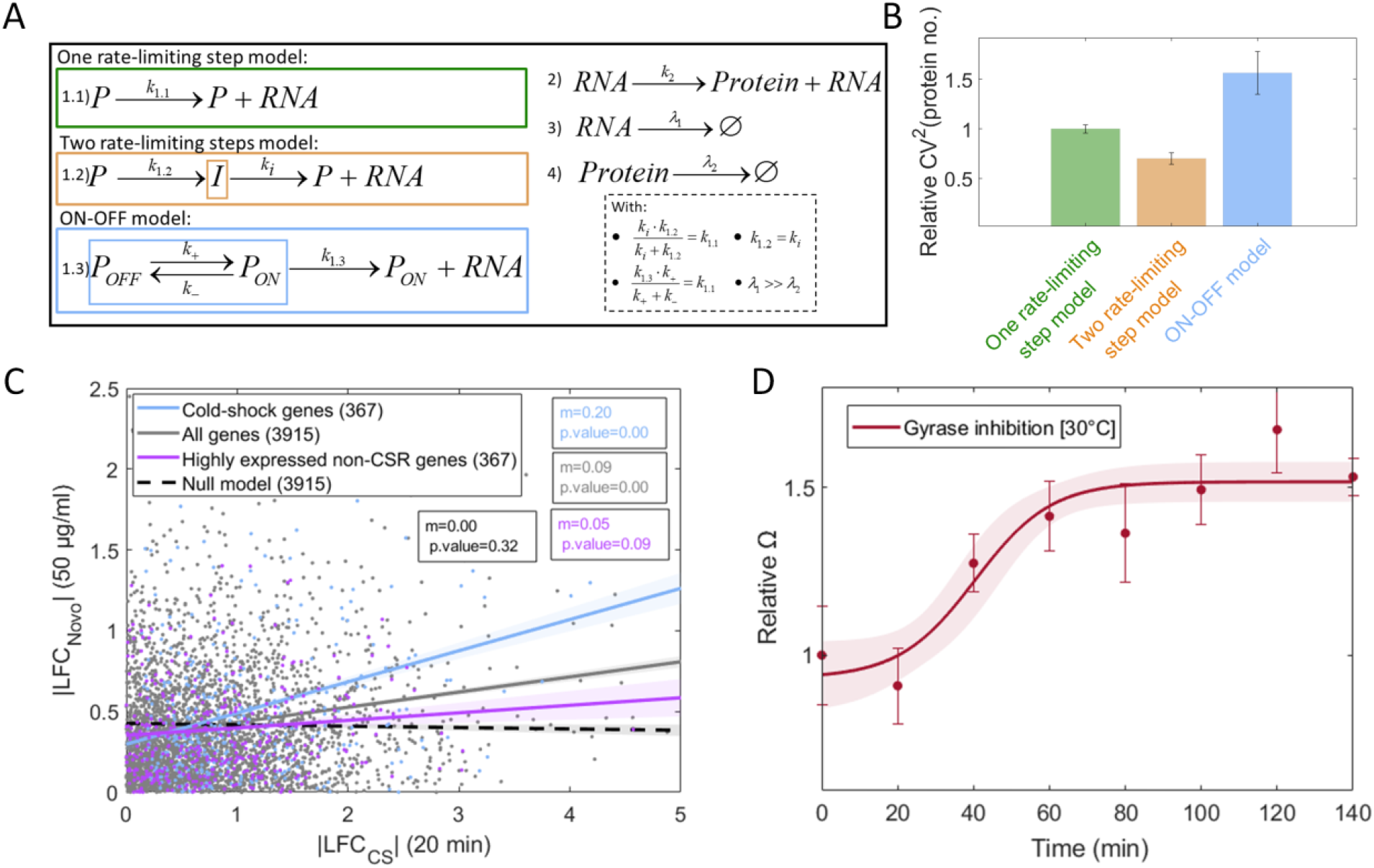
Nature of the short-term cold shock responses. **(A)** Three models were considered, differing in the reaction of transcription (reaction 1.1 for a one rate-limiting step model, reactions 1.2 for a two rate-limiting steps model, and reactions 1.3 for an ON-OFF model). All models include the same reactions for translation and RNA and protein decay (reactions 2, 3, and 4, respectively). The inset shows the conditions that the rate constants of the models must respect to impose identical mean protein numbers between them. **(B)** CV^2^ of protein numbers (relative to the one step model), as predicted by each model, assuming the parameter values in Table S8. Vertical error bars are the SEM. **(C)** Scatter plots of |LFC_NOVO_| after subjecting cells to 50 μg/mL Novobiocin (relative to a control condition, absent of Novobiocin) versus the |LFC_CS_| of mRNAs 20 min after shifting to 15°C (Methods Section 2.1). The Temperature RNAseq data has 4328 genes, while the Novobiocin dataset has 3948 genes. Common to both datasets are 3915 genes (grey circles). From the 3915 genes found, the blue circles correspond to 367 CSR. To the data, we fitted by OLS the best fit line (grey line and blue line, respectively). As a null model, we randomized the |LFC| for grey circles and fit by OLS the best-fit line between the correlation of the random pairs (dashed grey line). For each fit, we performed a likelihood ratio test between the zero-order polynomial and higher-order polynomials. P-values < 0.05 reject the null hypothesis that the best fit line is a horizontal line. **(D)** Effects of Novobiocin over time on Ω, which is best fit by a sigmoid function 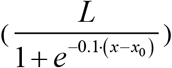, where *L* is the curve’s maximum value and *x*_*0*_ is the value of *x* of the midpoint of the sigmoid function. The red curve corresponds to the best-fit curve (that maximize R^2^). Measurements are performed by flow cytometer every 20 min, for 140 min. For each time point, we fit the function CV^2^=Ω/P [Bar-Even et al., 2006; Taniguchi et al., 2010].

First, we observed that the selection of this cohort occurs quickly, at the transcription level (Results Section 3.2), which excludes changes in k_2_ as the main cause for the selection. In support, we observed that changes in RNA and protein numbers in CS are correlated (Supplementary Figure S11B) and thus, no particularly relevant regulation is expected to be occurring during translation. Finally, we failed to find any statistically significant differences in the RBS sequence of the RNAs coded by the CSR genes and randomly selected genes (Supplementary Figure S8, see also Supplementary Section VIII), in what regards their Shine-Dalgarno (Table S9) as well as their start codon sequences (Table S10).

Second, RNA degradation in *E. coli* does not correlate with RNA sequence, abundance, or metabolic function [Bernstein et al., 2002; Chen et al., 2015; Deutcher et al., 2006], nor with cell responses to acute events [Bernstein et al., 2002]. Thus, we do not expect that changes in λ_1_ of CSR genes contributes to their selective responsiveness to CS.

We thus hypothesized that another mechanism, not present in the one-step model, ought to be responsible for the selective responsiveness of CSR genes, which includes the non-linear shift in the relationship between mean and noise (Figure 4B). We thus considered the emergence of an additional rate-limiting step in transcription initiation. A similar phenomenon has been reported [Buc and McClure, 1985] to occur on a synthetic promoter when shifting to temperatures lower than 20 °C. Also, it has been reported that tuning the supercoiled state of the DNA template can oppose this effect [Buc and McClure, 1985].

If the origin of the reduction in transcription rates is supercoiling buildup, then it can be accounted for by an ON-OFF process (e.g., by replacing reaction 1.1 by reactions 1.3, Figure 6A) [Chong et al., 2014; Palma et al., 2020; Oliveira et al., 2016]. Else, if the reduction results from the slowdown of the forward kinetics of transcription initiation, e.g., due to an isomerization process preceding open complex formation (Buc and McClure, 1985), then it can be modeled by two forward, rate-limiting steps in RNA production (e.g., by replacing reaction 1.1 by reactions 1.2, Figure 6A).

To determine which model is more realistic, we estimated their noise (CV^2^) for the same mean expression level. In detail, we tuned the three models so that they match in mean expression, by enforcing the relationships shown in the inset of Figure 6A between the rate constants (specific parameter values in Supplementary Table S8). Next, we performed stochastic simulations (Methods Section 2.4) and found that the ON-OFF model is the only one with higher noise than the one-step model and, thus, fits better the increase in Ω, at low temperatures (Figure 6B) meaning that, in CS, CV^2^ is higher than at optimal temperatures (for the same mean expression levels).

### 3.7 Response strength to cold shock is correlated with Positive Supercoiling sensitivity

We next explored the hypothesis that short-term responses to CS emerge from positive supercoiling sensitivity. For this, we performed RNA-seq after subjecting cells to 50 μg/mL Novobiocin, which inhibits Gyrase [Gellert et al., 1976; Mizuuchi et al., 1978] (Methods Section 2.1) and, thus, would cause a similar effect as CS if the hypothesis holds true.

From Figure 6C, the response strengths of CSR genes to Novobiocin are positively correlated to their response strengths to CS (blue balls in Figure 6C, p-value < 0.05), which supports the hypothesis. Further, CSR genes are more sensitive to Novobiocin than the average gene, i.e., have stronger responses, which further supports that they are more PSS.

On the other hand, it could instead be because their original expression in the control condition was relatively high, when compared to the average gene. To test this, we compared the response strength to Novobiocin of genes that are *not* CSR but, have similar expression levels in optimal conditions. We selected random cohorts with the same number of genes as the CSR cohort and the same average expression level in the optimal condition. We found that the best fitting line between their responses to CS and Novobiocin (Figure 6C) has a smaller slope than the line for CSR genes (and the two slopes can be statistically distinguished). We conclude that it is not their high expression level that explains why CSR genes are also PSS.

Given the above, we hypothesized that PSS is a key underlying mechanism of the short-term transcriptional program of cold shock responsiveness. To find if CSR genes are also PSS, we considered the genes whose responses to CS were stronger. To select them, we set a threshold between weak and strong at |LFC_CS_| = 0.8 (Supplementary Figure S19) since, below it, several p-values are close to the significance level (Supplementary Section I).

Next, to investigate if genes with strong CSR also have high PSS, we also needed to classify genes as having ‘high’ |LFC_NOVO_|. For this, we considered that the inclination of the best fitting lines in Figure 6C likely differ with perturbation strengths (e.g., adding more than 50 μg/mL Novobiocin would cause stronger LFCs [Palma et al., 2020]). Since, on average, the response strength of CSR genes to 15 °C was twice as strong as their response to 50 μg/mL Novobiocin, we classified responses of |LFC_NOVO_| > 0.4 as ‘strong’ (if their p-value < 0.05). Given this, 1215 out of 3948 genes of *E. coli* (∼31%) were classified as having a strong response to Novobiocin.

Given the classifications, of the 381 genes classified as CSR, 201 are strongly responsive to CS. Of these, we considered only 190, since the other 11 failed to obey the filtering criteria *iii*, in step I.b. in Supplementary Section I. Of the 190, 92 are strongly responsive to Novobiocin. Thus, approximately 50% of the CSR genes are also PSS, which is higher than expected by random chance.

Given this, we hypothesized that high sensitivity to PSB is at the core of the short-term, cold shock responsive transcriptional program of *E. coli*. Nevertheless, we also conclude that PSB is not the only means by which genes can be part of the cohort of quickly repressed genes during cold shocks. Finally, it is worth noting that, 1215 out of 3948 genes of *E. coli* exhibited |LFC_NOVO_| > 0.4 (p-value < 0.05), but only 92 of them were CSR. Thus, being PSS is not sufficient to be strongly, short-term CSR.

Given this, from *in vivo* single-cell, time lapse protein data (Methods section 2.1), we studied the dynamics of the 6 genes used to produce Figure 4B and investigated if their CS responsiveness is due to their PSS. In detail, if during CS, a rate-limiting step emerges in their dynamics (reaction 1.3 in the ON-OFF model in Figure 6A), we expect that the noise for a given mean expression level should be higher than during optimal conditions. For this, we further measured 4 additional CSR genes because they, in addition to pepN and ndk of the 6 genes, are the only ones out of the 381 CSR genes that: i) do not have any known input TFs and, thus, even in optimal conditions, should be less influenced by the TF network of *E. coli*; ii) their expression levels in control conditions are above background noise, and; iii) they are not integrated in a position of an operon structure other than the first one downstream the promoter.

Results in Supplementary Figure S18 show that, in accordance with the predictions, there is a decrease in mean protein levels during CS and Gyrase inhibition. Only two genes, pepN and feoA, exhibit increased levels, contrary to the model, after 60 and 100 mins following the addition of Novobiocin, respectively. This is, potentially, due to mid- and long-term phenomena (also respectively) occurring as part of the cellular response program to cold. For example, feoA has 4 input TFs, while pepN is closely spaced to another gene, ssuB, in a convergent configuration. Also, ssuB has no transcription termination site. As such, it can perturb pepN’s expression, e.g., by first repressing and then stopping doing so, when under the effects of Novobiocin.

Meanwhile, the overall ratio between noise and mean following Gyrase inhibition (Figure 6D) fits well a sigmoid, as it did when subjecting cells to CS (Figure 4B). The main difference between Figures 4B and 6D is that it takes less 20 min for the shift to occur following Novobiocin addition. This might be due to the slowing down of metabolic events during cold.

Finally, we note that the similarity in the mean changes in Ω is not used as criteria to support that the underlying mechanism is the same, because we tuned the Novobiocin levels to make the average strong of the perturbations similar.

### 3.8 The overlap between Gyrase and Nucleoid during CS increases

If the ON-OFF Model (Figure 6A) is in accordance with the process of gene expression during CS and, if Gyrases are responsible for removing promoters from their OFF state due to PSB [Chong et al, 2014], one would expect Gyrases to be more present in the DNA region during CS. Instead, if it was the two-step model (Figure 6A) that best described CS effects, then the RNAP would take longer to complete transcription initiation events and thus, would spend longer time at the DNA region.

To verify this, we measured by microscopy (Supplementary section III), prior to and during CS, the cell areas occupied by GyrA and RpoB, respectively. We also assessed how these areas overlapped with the area occupied by the nucleoid, observed by DAPI staining (Methods section 2.2), since temperature is known to perturb chromosomal DNA compaction [Goldstein et al., 1984; López-García et al., 2000].

Soon after cells enter CS, the nucleoid area decreases for the entire period of our gene expression measurements (Figure 7A). Since past studies observed a similar phenomenon when inhibiting Gyrase [Palma et al., 2020] and since the nucleoid area is a good proxy for nucleoid density [Gray et al., 2019], whose increase is a common effect of PSB [Eriksson et al., 2002], we hypothesize that Gyrase duty cycles increase during CS. In agreement, the overlap between the Gyrase and nucleoid regions increase during CS (Figure 7D, see example cells), even though the ‘Gyrase region’ decreased in size relative to the nucleoid region (Figure 7B.1 and B.2). Contrary to this, the overlap between the RNAP and nucleoid region does not increase during CS (Figure 7E, see example cell), even though the ‘RNAP region’ increased in size relative to the nucleoid region (Figure 7C.1 and C.2) which is in line with reduced number of available promoters for transcription initiation.

**Figure 7.**
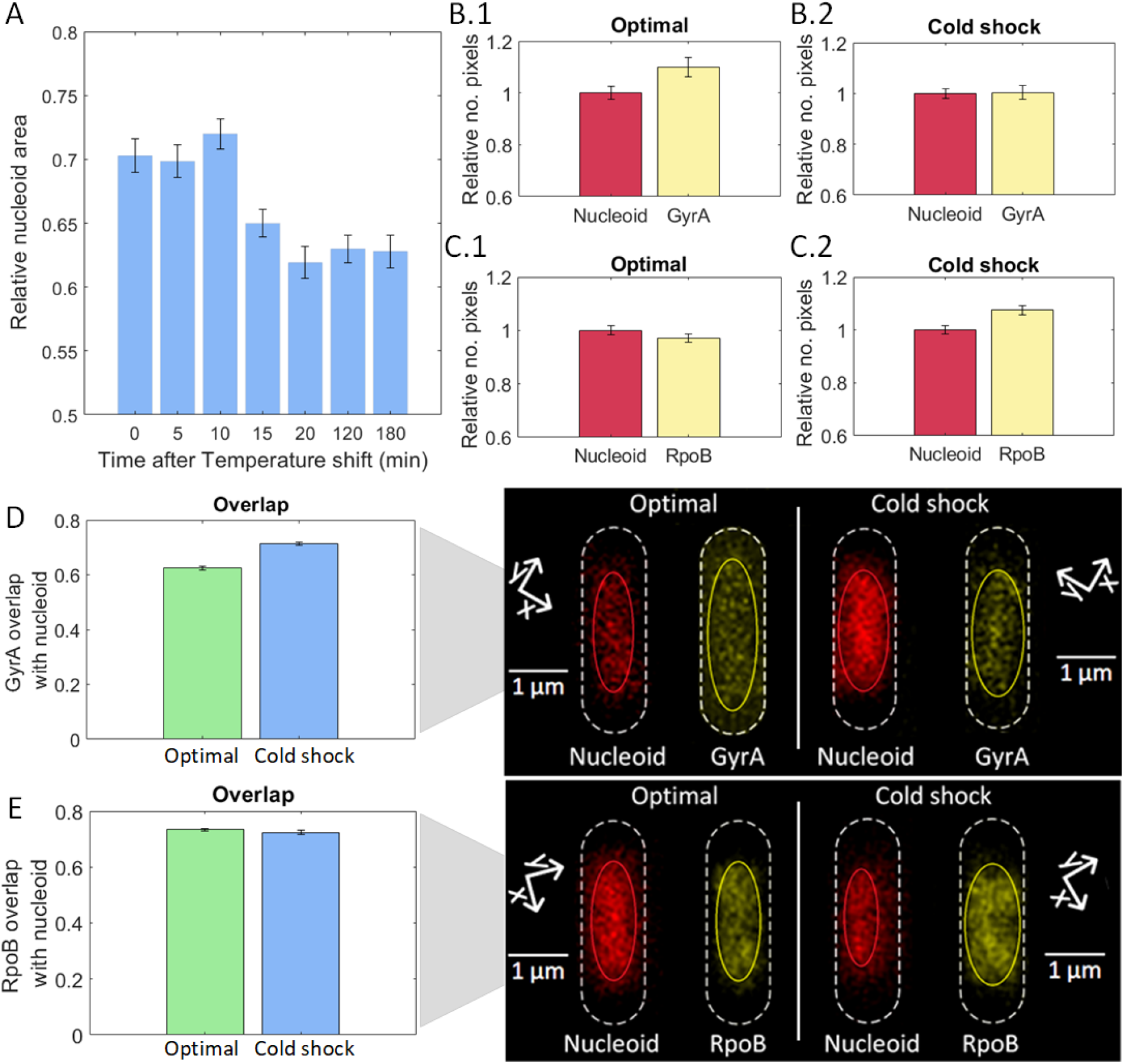
Model Study of biophysical parameters affected by cold shock; **(A)** Nucleoid area relative to cell area over time following CS, measured using ‘SCIP’ [Martins et al., 2018]. More than 500 cells analyzed per condition. **(B.1)** Size of YFP tagged GyrA relative to nucleoid size in optimal temperature. **(B.2)** Size of YFP tagged GyrA relative to nucleoid size after cold shock. **(C.1)** Size of YFP tagged RpoB (px) relative to nucleoid size (px) in optimal temperature. **(C.2)** Size of YFP tagged RpoB relative to nucleoid size after cold shock. **(D)** Relative overlapping between GyrA and the nucleoid in optimal and cold shock temperatures (Relative overlap is calculated as 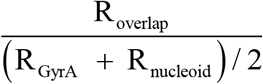, where R is “size of the region occupied by”). **(E)** Relative overlapping between the RpoB and nucleoid in optimal and cold shock temperatures. Also shown, example microscopy images of the same cell, with the nucleoid stained with DAPI and the GyrA/RpoB proteins tagged with YFP in optimal and cold shock temperatures, respectively. More than 400 cells analyzed per condition. Vertical error bars correspond to the standard error of the mean. Size is defined by number of pixels.

Overall, these results support the hypothesis that the ON-OFF model explains the underlying mechanism of a large number of short-term CS responsive genes.

### 3.9 Cellular energy levels decrease during cold shock

Gyrase ability to remove supercoils is ATP dependent [Rovinskiy et al., 2012; Gubaev et al., 2016]. Also, Gyrase numbers did not increase during CS (Figure 2D). As such, a decrease in ATP levels could contribute to make PSB an ‘efficient’ mechanism underlying CSR genes. To study this, we measured ATP levels in the control and CS conditions (Methods section 2.3). These levels quickly decrease during CS (Supplementary Figure S13). This furthers supports the ON-OFF Model, as it suggests that the average time to escape OFF states increases in CS.

### 3.10 Relative Ω as a function of OFF-ON rates

From above, we expect CS to alter how Ω is regulated due to the emergence of an ON-OFF step controlling transcription. We estimated the expected ratio between values of Ω at cold and optimal conditions assuming ON-OFF and one-step models (Figure 6A), respectively (Supplementary Section V.e.). From there:

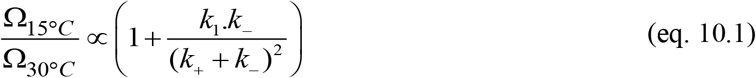

Equation 10.1 informs on how the ratio Ω_15°*C*_ / Ω_30°*C*_ is expected to be affected by the rate constants controlling the ON-OFF steps, k_+_ and k_-_ (Reactions 1.3 in Figure 6A), and the transcription activity from active promoters, k_1_.

We do not expect k_1_ to be a major regulator of this ratio, since this rate constant is present in the one-step model, which was unable to mimic the measurements. Meanwhile, of the two remaining events controlling promoter activity (Reactions 1.3, Figure 6A) only promoter escape from the OFF state (regulated by *k*_+_) is energy consuming [Gubaev et al., 2016]. Thus, this event is expected to be most decelerated one during CS. We therefore hypothesized that k_+_ is the most temperature sensitive parameter in equation 10.1.

We therefore investigated the relationship between k_+_ and temperature. We explored four temperature sensitive models of k_+_ which were fitted to the empirical data from Figure 4B. Models and best fitting parameter values are shown in Supplementary Table S12, while results of the fitting are shown in Supplementary Figure S12. From the R^2^ values, the best fitting model assumes that k_+_ changes over time following an exponential function.

## 4. Discussion

We identified a large number of short-term CSR genes and studied what causes their quick repression in CS. A few of them are likely responsive due to being in an operon with upstream CSR genes, but the majority appear to be independently responsive to CS. Interestingly, following CS, CSR genes rapidly decrease expression level, while their noise relative to the mean expression increases. This increase in noise is consistent with the emergence of transient locking events of transcription. Since a similar phenomenon was observed following Gyrase inhibition [Chong et al., 2014; Palma et al., 2020] and because we observed here that a large number of CSR genes is also highly sensitive to PSB, we hypothesized that the responsiveness of a large number of CSR genes emerges from their PSB responsiveness. Meanwhile, we observed that Gyrase converges to the nucleoid and that cell energy decreases during CS, suggesting that the number of promoters locked due to PSB increases during CS. We therefore proposed a model of the responsiveness of CSR genes based on their temperature sensitive PSB.

To our knowledge, temperature sensitive PSB is the first identified physical mechanism of how an *E. coli* gene can be CS responsive, and evidence suggests that it is present in nearly half of *E. coli*’s CSR genes. This finding, first hypothesized in [Oliveira et al., 2019], opens an avenue for the engineering of future synthetic, temperature sensitive and temperature resistant gene regulatory circuits, whose functioning could be tuned by the adaptive regulation of Gyrase activity. Further, we expect that it will contribute to learning how the short- and long-term transcriptional programs of *E. coli* responsive to CS have evolved.

Nevertheless, we found that not all short-term CSR genes (∼ 50% of them) are responsive to PSB. As such, there must be other mechanisms by which genes become short-term CSR. Similarly, we also observed that not all genes responsive to PSB are also short-term CSR, implying that being responsive to PSB does not suffice to be CSR. Finally, it also remains unclear how PSB percolates genes in the same operon in a manner that, while some genes downstream a CSR gene are also CSR, many are not.

Given this, much study is still needed to identify the various possible set of features that can make a gene CSR. Potentially, genes could be CSR by CS-based locking of their rate-limiting steps during transcription initiation, as reported for a synthetic promoter [Buc and McClure, 1985]. This could explain how, in 4 out of the 30 genes measured at the protein level, noise did not increase although the mean levels decreased. However, we do not expect this to be common in most CSR genes, since the size of the cellular region occupied by RNAP during CS increased. Finally, it may be that, in some genes, their RNA or proteins have increased decay rates during CS, rather than altered production rates.

Bacterial transcriptional programs of cold shock responsiveness are critical survival skills that affect a wide range of vital Human activities [Phadtare et al., 2004]. It should be possible to perturb this ability by introducing it in the cells’ synthetic circuits, with a temperature sensitivity based on the PSS of its component genes. By interfering with CSR genes that rely on PSB, from our results, we expect that half of the transcriptional program of short-term response would be affected. As such, this is a viable strategy with potentially great rewards. For example, if we could activate natural bacterial CS transcriptional programs in the absence of cold, we would be able to slow down infections and food spoilage. We would also be able to enhance natural bacterial programs in low-temperature bioreactors (e.g., responsible for fermentation in the dairy industry), making them more cost-efficient. Meanwhile, if we could deactivate it when desired, we would be able to enhance bio fertilization and plant resistance to bacteria, among others.

## Supplementary data

See supplementary file attached.

## Author Contributions

A.S.R., C.S.D.P., and S.D. conceived the study and ASR supervised it. S.D. planned and executed the measurements, to which C.S.D.P. and V.C. contributed to. C.S.D.P. planned and executed the data analysis to which S.D., I.S.C.B., M.N.M.B., B.L.B.A. and R.J. contributed to. A.S.R., S.D., and

C.S.D.P. drafted all documents, which were revised by all co-authors. S.D. and C.S.D.P. contributed equally and have the right to list their name first.

## Funding

This work was supported by the Jane and Aatos Erkko Foundation [10-10524-38 to A.S.R.]; Suomalainen Tiedeakatemia to C.S.D.P.; Finnish Cultural Foundation [00200193 and 00212591 to I.S.C.B., and 50201300 to S.D.]; EDUFI Fellowship [TM-19-11105 to S.D.]; Tampere University Graduate Program to V.C., M.N.M.B., and B.L.B.A.. The funders had no role in study design, data collection and analysis, decision to publish, or preparation of the manuscript.

## Conflict of Interest

The authors have no competing interests.

## Notes

### Competing Interest Statement

The authors have declared no competing interest.

