## Supplementary Information for "Positive supercoiling buildup is a trigger of *E. coli*’s short-term response to cold shock"

**Supplementary Methods**

### I. RNA-Seq experiments and analysis

#### *I.a) Sample preparation*

We shifted the temperature for cells in the mid-exponential growth phase (OD=0.3) to 15°C (here named minute 0) and collected cells from three independent colonies, as well as after 20, 80 and, 180 minutes, respectively. The same was done with cell colonies not subjected to the temperature shift, for purposes of control. Meanwhile, when subjecting cells to the antibiotic Novobiocin (50 μg/mL), the cells were only collected 120 minutes after the treatment, to reduce cell-to-cell diversity due to e.g., different absorption times, as in [Chong et al., 2014].

After collecting the samples, 5 ml of the culture was immediately treated with a double volume (10 mL) of RNA protect bacteria reagent (Qiagen, Germany) for 5 minutes at room temperature, to prevent RNA degradation. Next, treated cells were pelleted and frozen at -80°C overnight. The next morning, total RNA was extracted using the RNeasy kit (Qiagen, Germany).

#### *I.b) Sequencing*

Extracted RNA was treated twice with DNase (Turbo DNA-free kit, Ambion, USA) and quantified using Qubit 2.0 Fluorometer RNA assay (Invitrogen, Carlsbad, CA, USA). The total RNA quality was determined using a 1% agarose gel stained with SYBR Safe (Invitrogen, Carlsbad, CA, USA), where RNA was detected using UV in a Chemidoc XRS imager (Biorad, USA). RNA integrity was measured by the Agilent 4200 TapeStation (Agilent Technologies, Palo Alto, CA, USA).

RNA library preparations, sequencing, and quality control analysis of sequenced data were conducted at GENEWIZ, Inc. (Leipzig, Germany). In detail, ribosomal RNA depletion was performed using Ribo-Zero Gold Kits (Bacteria probe) (Illumina, San Diego, CA, USA), while the RNA sequencing library was prepared using the NEBNext Ultra RNA Library Prep Kit.

The sequencing libraries were multiplexed and clustered on one lane of a Flowcell, which was loaded on an Illumina HiSeq 4000 instrument (after cold shock) or on an Illumina NovaSeq 6000 instrument (after Gyrase inhibition). In both instruments, the samples were sequenced using a single-index 2x150 Paired-End (PE) configuration. Image analysis and base calling were conducted by the HiSeq Control Software (Illumina HiSeq) and by the NovaSeq Control Software v1.7 (Illumina NovaSeq). The raw sequence data (.bcl files) was converted into "fastq" files and de-multiplexed using Illumina bsl2fastq v.2.20. One mismatch was allowed for index sequence identification.

#### *I.c) RNA-seq data analysis pipeline*

i) RNA sequencing reads were trimmed to remove possible adapter sequences and nucleotides with poor quality using Trimmomatic [Bolger et al., 2014] v.0.36. ii) Trimmed reads were mapped to the reference genome, *E. coli* MG1655 (NC_000913.3), using the STAR aligner v.2.5.2b (after cold shock) or the Bowtie2 aligner v.2.3.5.1 (after Gyrase inhibition), generating BAM files [Dobin et al., 2013; Langmead et al., 2012]. iii) Unique gene hit counts were calculated with 'featureCounts' from the Rsubread R package (v.1.34.7) [Liao et al., 2019]. Genes with less than 5 counts in more than 3 samples, and genes whose mean counts were smaller than 10 were removed from further analysis. iv) The read counts were used for downstream differential expression analysis. The DESeq2 R package (v.1.24.0) [Love, et al., 2014] was used to calculate log2 of fold changes (LFC) of RNA levels between group of samples and calculate p-values using Wald tests (function 'nbinomWaldTest'). We also calculated relative abundances of mRNA in a given condition using the Transcripts Per Million (TPM) normalization [Li et al., 2011].

Finally, the sequencing platforms for the cold shock experiment and for the Novobiocin experiment differ for logistical reasons (similarly, the aligners differ). Consequently, we avoided quantitative comparisons between genes responses (e.g., between specific numbers). Our conclusions are based on qualitative comparisons (i.e., we checked if the changes in the two perturbations are linearly correlated or not).

**II. Flow cytometry and data analysis**

We measured cells fluorescence using an ACEA NovoCyte Flow Cytometer (ACEA Biosciences Inc., San Diego, USA). Cells were diluted (1:10000) into 1 mL of Phosphate buffer saline (PBS) solution, vortexed for 10 seconds. In each condition, 3 biological replicates were obtained. In each replicate, we collected data from 50 000 cells. The flow rate was set to 14 µL/minute. The data was collected by the Novo Express software (ACEA Biosciences Inc.).

For detecting YFP, we used a blue laser (488 nm) for excitation and the fluorescein isothiocyanate detection channel (FITC-H) (530/30 nm filter) for emission, with a core diameter of 7.7 µM and a PMT voltage of 600. For detecting mCherry, we used the PE-Texas Red fluorescence detection channel (615/20 nm) for emission, with a PMT voltage of 584.

The lower bound for the detection threshold in FSC-H was set to 5000 to remove interference from particles. We also removed the 1% highest FITC-H values. Further, to remove abnormal cells, we then used an iterative procedure to discard outliers, which are those points whose vertical distance from the best-fit function is larger than 1 [Bar-Even et al., 2006]. The process always converged in 1 to 2 iterations. Finally, we searched for additional abnormal measurements at the single gene level in the 3 repeats, but we did not find any.

### III. Microscopy and image analysis

Cells were pelleted and re-suspended in ~100 µL of the remaining media. Three microliters of cell suspension were placed on a 2% agarose gel pad made up of M9 medium and kept in between the round microscope slide and a coverslip. It took ~5 min to move cells from the incubator to the microscope and start the observation. This includes the assembly of the microscope imaging chamber containing the slides and cells. Cells were visualized by confocal microscopy with a 100x objective. Phase-contrast images were taken by an external phase-contrast system. YFP tagged strain was visualized by a 488 nm laser and a 514/30 emission filter, while DAPI stained nucleoids were visualized by a 405 nm laser and a 447/60 emission filter. Phase contrast and confocal images were taken simultaneously.

Analysis of the microscopy images was performed using the CellAging software [Häkkinen et al., 2013], for segmenting cells from phase contrast images, and the SCIP software [Martins et al., 2018] to segment nucleoids and characterize the spatial distribution of fluorescently (YFP) tagged GyrA and RpoB (both from the YFP fusion library).

### IV. Correction for cellular auto-fluorescence in Flow Cytometry data

When assessing the single-cell distributions of protein expression levels measured by flow-cytometry, we corrected for the cell auto-fluorescence [Bahrudeen et al., 2019; Galbusera et al., 2020]. For this, we first measured by flow-cytometry the auto-fluorescence of control cells (i.e., absent of YFP fusions). Next, we corrected the mean fluorescence measured by flow cytometer by applying equation IV.1 [Galbusera et al., 2020]:

$$M_p = M_T - M_{cell} \tag{IV.1}$$

Here, $M_p$ is the mean cell fluorescence due to YFP presence alone, after subtracting the cell auto-fluorescence. Meanwhile, $M_T$ is the mean cell fluorescence measured by flow cytometry, while $M_{cell}$ is the mean cell auto-fluorescence. Similarly, to correct the variance $\sigma^2$, we apply equation IV.2 [Galbusera et al., 2020]:

$$\sigma_p^2 = \sigma_T^2 - \sigma_{cell}^2 \tag{IV.2}$$

From equations IV.1 and IV.2 one can derive equation IV.3, to estimate the corrected squared coefficient of variation, $CV_p^2$, of the single-cell distribution of protein expression levels:

$$CV_p^2 = \left( \frac{\sigma_p}{M_p} \right)^2 \qquad \text{(IV.3)}$$

For correcting the skewness (S) of the distribution we apply equation (IV.4) as in [Bahrudeen et al., 2019]:

$$S_p = \frac{S_T \cdot \sigma_T{}^3 - S_{cell} \cdot \sigma_{cell}{}^3}{\sigma_p{}^3} \qquad \text{(IV.4)}$$

**V. Derivations**

**Va. Squared coefficient of variation and skewness assuming a Γ distribution of single-cell protein numbers**

From [Taniguchi et al., 2010], most single-cell distributions of protein numbers in E. coli are well described by a Γ distribution. If that holds, the first three moments of single-cell distributions of protein numbers should be given by:

$$M = k\theta \qquad \text{(Va.1)}$$

$$\sigma^2 = k\theta^2 \qquad \text{(Va.2)}$$

$$S = \frac{2}{\sqrt{k}} \qquad \text{(Va.3)}$$

where M, $\sigma^2$, S, k and θ are the mean, variance, skewness, shape parameter, and scale parameter of a Γ distribution, respectively. From equations V.1 and V.2:

$$CV^2 = \frac{\sigma^2}{M^2} = \frac{\theta}{M} \qquad \text{(Va.4)}$$

This relationship was empirically validated in [Bar-Even et al., 2006][ Taniguchi et al., 2010]. Finally, from equation V.1 and V.3:

$$S = \frac{2}{\sqrt{k}} = \frac{2}{\sqrt{\frac{M}{\theta}}} = \frac{2}{\sqrt{M}} \cdot \sqrt{\theta} \qquad \text{(Va.5)}$$

**V.b Derivation of Ω assuming the 1-step model**

The steady state solutions for mean RNA and protein numbers, assuming the one step Model (reactions 1.1, 2, 3, and 4 in Figure 6), are given by, respectively [Taniguchi et al., 2010]:

$$M_{RNA} = \frac{k_{1.1}}{\lambda_1} \tag{Vb.1}$$

$$M_P = \frac{M_{RNA} \cdot k_2}{\lambda_2} = \frac{k_{1.1} \cdot k_2}{\lambda_1 \cdot \lambda_2} \tag{Vb.2}$$

Meanwhile, the variance of the single-cell protein numbers is given by [Taniguchi et al., 2010]:

$$\sigma_P^2 = \frac{k_{1.1} \cdot k_2}{\lambda_1 \cdot \lambda_2} \cdot \left(1 + \frac{k_2}{\lambda_1 + \lambda_2}\right) \tag{Vb.3}$$

From (Vb.2) and (Vb.3):

$$CV_P^2 = \frac{y_1 \cdot y_2}{k_{1.1} \cdot k_2} \cdot \left(1 + \frac{k_2}{\lambda_1 + \lambda_2}\right) = \frac{1}{M_P} \cdot \left(1 + \frac{k_2}{\lambda_1 + \lambda_2}\right) \tag{Vb.4}$$

Here, we define the constant Ω as:

$$\Omega = \left(1 + \frac{k_2}{\lambda_1 + \lambda_2}\right) \tag{Vb.6}$$

This result is in line with past results in [Bar-Even et al., 2006]. Also, given (Va.4), we expect θ = Ω. As such, we refer to this constant as Ω.

**V.c Derivation of the squared coefficient of variation assuming the 1-step model**

Assuming Model 1.1 in Figure 6, from (Vb.2), (Vb.4), and (V.5):

$$CV_p^{\,2} = \frac{1}{\frac{k_1 \cdot k_2}{\lambda_1 \cdot \lambda_2}} \cdot \left(1 + \frac{k_2}{\lambda_1 + \lambda_2}\right) \tag{Vc.1}$$

**V.d Derivation of the Squared Coefficient of Variation assuming the ON-OFF model**

Assuming the ON-OFF model (Model 1.3 in Figure 6), the mean RNA and protein numbers in single cells at steady state are, respectively:

$$M_{RNA} = \frac{k_+}{k_+ + k_-} \cdot \frac{k_{1.3}}{\lambda_1} \tag{Vd.1}$$

$$M_P = \frac{M_{RNA} \cdot k_2}{\lambda_2} = \frac{k_+}{k_+ + k_-} \cdot \frac{k_{1.3} \cdot k_2}{\lambda_1 \cdot \lambda_2} \tag{Vd.2}$$

Meanwhile, the variance is [Taniguchi et al., 2010]:

$$\sigma_P^2 = \frac{k_+}{k_+ + k_-} \cdot \frac{k_{1.3} \cdot k_2}{\lambda_1 \cdot \lambda_2} \cdot \left( 1 + \frac{k_2}{\lambda_1 + \lambda_2} \left( 1 + \left( 1 - \frac{k_+}{k_+ + k_-} \right) \frac{k_{1.3}(k_+ + k_- + \lambda_1 + \lambda_2)}{(k_+ + k_- + \lambda_1)(k_+ + k_- + \lambda_2)} \right) \right) \quad \text{(Vd.3)}$$

From equations Vd.3 and Vd.3:

$$CV_P^2 = \frac{1}{M_P} \cdot \left( 1 + \frac{k_2}{\lambda_1 + \lambda_2} \left( 1 + \left( 1 - \frac{k_+}{k_+ + k_-} \right) \frac{k_{1.3}(k_+ + k_- + \lambda_1 + \lambda_2)}{(k_+ + k_- + \lambda_1)(k_+ + k_- + \lambda_2)} \right) \right) \quad \text{(Vd.4)}$$

From Vd.4 and Va.4, Ω should equal:

$$\Omega = \left( 1 + \frac{k_2}{\lambda_1 + \lambda_2} \left( 1 + \left( 1 - \frac{k_+}{k_+ + k_-} \right) \frac{k_{1.3}(k_+ + k_- + \lambda_1 + \lambda_2)}{(k_+ + k_- + \lambda_1)(k_+ + k_- + \lambda_2)} \right) \right) \quad \text{(Vd.5)}$$

#### V.e Changes in Ω due to cold shock

To estimate the change in Ω due to shifting to CS we considered that, prior to cold shock, the CS responsive genes due to sensitivity to PSB are not significantly affected by locking due to PSB (since they are relatively highly expressing in optimal growth conditions). As such, their dynamics in optimal conditions should be well modeled by Model 1.1 in Fig 6A in the main manuscript. On the other hand, when subject to CS, we expect that frequent locking due to PSB will be the main responsible for their negative response. Thus, the appropriate model during CS should be Model 1.3 in Fig 6A in the main manuscript.

Thus, given the results in supplementary sections V.b and V.d, the change in Ω when shifting to cold shock should equal:

$$\frac{\Omega_{15°C}}{\Omega_{30°C}} = \frac{\left( 1 + \frac{k_2(15°C)}{\lambda_1(15°C) + \lambda_2(15°C)} \left( 1 + \left( 1 - \frac{k_+}{k_+ + k_-} \right) \frac{k_1((k_+ + k_-) + \lambda_1(15°C) + \lambda_2(15°C))}{((k_+ + k_-) + \lambda_1(15°C))((k_+ + k_-) + \lambda_2(15°C))} \right) \right)}{\left( 1 + \frac{k_2(30°C)}{\lambda_1(30°C) + \lambda_2(30°C)} \right)} \quad \text{(Ve.1)}$$

Next, consider that, from Table S7, $\frac{k_2}{\lambda_1 + \lambda_2}$ = 14.9, which is much larger than 1. Thus, for simplicity, we replace $\left( \frac{k_2}{\lambda_1 + \lambda_2} + 1 \right)$ by $\frac{k_2}{\lambda_1 + \lambda_2}$. Further, we consider that, in *E. coli*, RNA degradation rates are much higher than protein degradation rates, i.e., $\lambda_1 >> \lambda_2$ (Table S7). Given this, we also replace $(\lambda_1 + \lambda_2)$ by $\lambda_1$. Consequently:

$$\frac{\Omega_{15^\circ C}}{\Omega_{30^\circ C}} = \frac{k_2(15^\circ C)\times\lambda_1(30^\circ C)}{k_2(30^\circ C)\times\lambda_1(15^\circ C)}.\left(1+\left(1-\frac{k_+}{k_+ + k_-}\right)\frac{k_1((k_+ + k_-)+\lambda_1(15^\circ C)+\lambda_2(15^\circ C))}{((k_+ + k_-)+\lambda_1(15^\circ C))((k_+ + k_-)+\lambda_2(15^\circ C))}\right) \quad \text{(Ve.4)}$$

This, by derivation, can be simplified to:

$$\Leftrightarrow \frac{\Omega_{15^\circ C}}{\Omega_{30^\circ C}} = \frac{k_2(15^\circ C)\times\lambda_1(30^\circ C)}{k_2(30^\circ C)\times\lambda_1(15^\circ C)}.\left(1+\left(\frac{k_1.k_-}{(k_+ + k_-)^2}\right)\right) \quad \text{(Ve.4)}$$

We note that we expect that the term $\left(\frac{k_1.k_-}{(k_+ + k_-)^2}\right)$ is the one containing the most temperature sensitive rate constants, given that model 1 in Figure 6 includes the other rates constants ($k_2$ and $\lambda_1$) and could not explain the dynamics following cold shock for the reasons listed in section 3.6 in the main manuscript.

**VI: Effects of cell division on Ω**

At temperatures above cold shock (30°C, 25°C and 20°C), the cells exhibited significant doubling times (Figure 2A). Meanwhile, at CS they did not divide (Figure 2A).

Cell division can increase single-cell variability in protein numbers, provided asymmetries in the partitioning of RNA and protein numbers between sister cells (for a review, see [Baptista et al., 2020]). This difference between the conditions, could affect the comparison of the contribution of noise in gene expression in optimal and CS conditions.

We thus estimated the effects of cell division on Ω at temperatures above cold shock (30°C, 25°C and 20°C) if the division rate at 30°C, 20°C and 25°C was null.

Since we are considering temperatures above cold shock (30°C, 25°C and 20°C), we assume the 1-step model (model 1.1 in Figure 6). Next, we assume that $\lambda_1 >> \lambda_2$, since, in general RNA degrades much faster than proteins [Taniguchi et al., 2010]. Given this, from Equation V.b.6 in supplementary section V.b:

$$\frac{\Omega_{30^\circ C(\lambda_d=7\times10^{-5})}}{\Omega_{30^\circ C(\lambda_d=0)}} = \frac{\left(1+\frac{k_2}{(\lambda_1+\lambda_d)+(\lambda_2+\lambda_d)}\right)}{\left(1+\frac{k_2}{\lambda_1+\lambda_2}\right)} = \frac{\left(\frac{k_2}{(\lambda_1+\lambda_d)+(\lambda_d)}\right)}{\left(\frac{k_2}{\lambda_1}\right)} = \frac{\lambda_1}{\lambda_1+2\lambda_d} \quad \text{(VI.1)}$$

Next, from [Bernstein et al., 2002], we assume that, on average $\lambda_1$ =0.004s$^{-1}$. Also, we measured the mean cell division rate at 30°C-20°C to be 241 min$^{-1}$ (Figure 2A). This inverse should correspond to the protein and RNA dilution rates due to cell division and it equals: $\lambda_d$ =7×10$^{-5}$ s$^{-1}$. As such:

$$\frac{\Omega_{30°C(\lambda_d=7\times10^{-5})}}{\Omega_{30°C(\lambda_d=0)}}=0.97 \qquad \text{(VI.1)}$$

Given this, if cells were not dividing in optimal conditions, Ω would be 3% higher. This is within the 90% confidence interval of the green line in Figure 4B. Thus, we do not include cell division in the models.

**VII: Promoter sequence logos**

Promoter sequence logos were created using WebLogo [Crooks et al., 2004]. From positions -25 to -1 of each promoter, it counts in how many promoters is each nucleotide present. Then, it piles up the nucleotides (A, C, T, G), sorted from the rarest in the bottom to the most frequent in the top.

The height of each nucleotide letter in the plot, in each position, equals the frequency multiplied by the total information at that position. That total information is quantified by the difference between the maximum uncertainty at any position ($\log_2$(n), where n = 4 is the number of possible codons) and the uncertainty given the frequencies found, also quantified by Shannon's information: $\log_2 n-\sum_{i=1}^{4} f_i \times \log_2(f_i)$. Given this, we expect that DNA with more conserved positions will have more 'bits' (Schneider and Stephens, 1990).

**VIII: RBS and start codon sequences of CSR genes**

RNA translation rates are controlled by the rate at which ribosomes are recruited to the ribosome binding site (RBS) region of the RNA, along with the rate at which they then initiate translation. The recruitment rate differs with the RBS sequence [Ringquist S et al., 1992] and the genome wide consensus sequence of RBSs is "5'-AGGAGG-3'", being named the Shine-Dalgarno (SD) sequence [Saito et al., 2020].

Meanwhile, the rate of translation initiation is influenced by the start codon upstream the RBS. In *E. coli*, 83% of the start codons have the sequence AUG (3542/4284), 14% (612) the sequence GUG, 3% (103) the sequence UUG [Blattner, 1977] and a couple the sequence AUU [Sacerdot et al., 1982; Missiakas et al.,1993].

We obtained the mean and standard deviation of the distributions of the p-distances (Supplementary Section X) of the RNAs coded by CSR genes to the SD (Table S9) and to each of the 4 start codons sequences (Table S10) and studied if they differ significantly from the mean and standard deviation of p-distances of the genome wide distribution. Finally, since the distance (in number of nucleotides) between the SD sequence and the start codon can affect translation initiation rates [Saito et al., 2020], we also compared them as above.

From Supplementary Figure S8, not only the consensus levels of the two cohorts are the same (bit values of 0.5 between positions -10 and -15 and 1 in the region -1 to -3), but the distances between them are, in both cohorts, 5 nucleotides.

To support, we also compared the sequence logos of the 25 nucleotides upstream of the start codons [Wenfa, 2019] (Supplementary Figure S9) of CSR genes and the genome wide distribution. Again, we find little differences between the logos in Fig. S9A and S9B.

**IX: Estimation of the average transcription rate of CSR genes during optimal growth conditions**

From [Taniguchi et al., 2010], the mean RNA numbers (as measured by FISH) of a CSR gene during optimal growth is 0.35 per cell. Given the 1-step model (model 1.1 in Figure 6), which is applied during optimal growth, the mean number of RNAs per cell in steady state is given by:

$$k_1 = M_{RNA} \times \lambda_1 \qquad \text{(VIII.1)}$$

Assuming $\lambda_1$=0.004 s$^{-1}$ [Bersntein et al., 2002; Selinger et al., 2003], $k_1$ is estimated to be $1.4\times10^{-3}$ s$^{-1}$.

**X: P-distances**

We calculated the p-distance between a promoter sequence and the consensus sequence (sequence composed of the most common nucleotide for each position of the sequence). The p-distance is the fraction of nucleotides of the promoter sequence that differ from the consensus sequence. Thus, it ranges from [0,1], where 0 corresponds to identical sequences and 1 to sequences whose nucleotides differ in every position. For genes with more than one promoter, we obtained the average of the p-distance of each promoter.

**XI: Correlation between RNA and protein numbers**

RNA and protein numbers are expected to be positively correlated in bacteria, since transcription and translation are mechanically bound [Dahan, O. et al., 2011; Yanofsky, C. et al., 1981; Proshkin, S. et al., 2010] and because most gene expression regulation occurs during transcription initiation [Alberts, B. et al., 2008].

To assess if this holds true during cold-shock, we searched for correlations between LFC's (Supplementary section I), as measured by RNA-seq at 20 and 80 min after the temperature shift, and the corresponding LFC's in protein numbers, measured by flow-cytometry (Supplementary section II) at 120 min and 180 min after the temperature shift. The lag of 100 minutes between RNA and protein measurements should suffice for changes in numbers of the former to propagate to the latter. The list of genes tested is shown in Table S1. Results in Figure S11B show that changes in RNA and protein numbers are correlated during CS.

**XII: Quantifying ATP using spectrophotometry**

To quantify the fluorescence from GFP tagged ATP inside cells, we use the method in [Yaginuma et al., 2014]. First, the total cell fluorescence at excitation wavelength λ is given by:

$$F_\lambda = F_\lambda^m + f_\lambda^{bg} \cdot C + f_\lambda^p . C \qquad \text{(XI.1)}$$

*F* stands for total fluorescence and *C* for the number of cells. Meanwhile, $F^m$ stands for media fluorescence, $f^{bg}$ for single-cell fluorescence background and $f^p$ for single-cell protein fluorescence (in our case, λ equals 400 nm in one case and 494 nm in the other).

Meanwhile, the total cell fluorescence (without ATP sensors) is:

$$F_\lambda^c = F_\lambda^m + f_\lambda^{bg} \cdot C \qquad \text{(XI.3)}$$

The subtraction of (XI.1) from (XI.3) corrects for media and cell background autofluorescence:

$$F_\lambda - F_\lambda^c = f_\lambda^p . C \qquad \text{(XI.5)}$$

Given this, the fluorescence from ATP-GFP from a cell is estimated by:

$$\frac{f_{\lambda=494}^p(t)}{f_{\lambda=400}^p(t)} = \frac{F_{\lambda=494}(t) - F_{\lambda=494}^c(t)}{F_{\lambda=400}(t) - F_{\lambda=400}^c(t)} \qquad \text{(XI.7)}$$

**XIII: Analysis of the AT and CG content of the promoters**

From Regulon DB, we obtained the lists of all 8791 promoters and of all 3700 transcription units (TUs) [Santos-Zavaleta et al., 2019]. We then filtered the promoter list to contain only the 2355 promoters associated to TUs, and subsequently discarded 93 promoters with unknown sequence. The resulting list was comprised of 2262 promoters, each with a sequence spanning from 60 nucleotides upstream the transcription start site (TSS) to 20 nucleotides downstream (i.e., from positions -60 to +20, with the TSS assumed to be in the position +1).

Next, for each promoter, we extracted the sequences from positions -60 to -35, positions -35 to -10, and positions -10 to +1 (the -35 to -10 being the sequence that most influence the RNAP binding [deHaseth et al., 1998], while the others were used for comparison). Similarly, we also extracted the same sequences of 443 CSR genes with known promoter sequences. Finally, for each set of sequences, we calculated the fractions of A, C, G and T. Finally, the AT (or GC) content of each promoter was calculated by summing the fractions of A and T (or C and G, respectively).

**XIV. Estimation of a lower bound of skewness**

Supplementary Figure S10, informs on a lower bound for noise ($CV^2 \sim 0.38$) since noise and mean are no longer correlated below that value. Based on this lower bound, we estimated a lower bound of skewness. From the equations in Table S6, one can write:

$$S = 2 \cdot CV \qquad \text{(XIII.1)}$$

We estimate a lower bound for the skewness to equal ~1,23, in agreement with the data (Figure 5), in that below that value, the predicted and empirical skewness do not correlate.

**XV. Null models and statistical tests**

In general, to create null-models of how variable X affects variable Y, we performed random sampling without replacement of both *X* and *Y* datapoints. The number of samplings and the sampling size (number of samples in each sampling) are set to the maximum array size allowed by MATLAB (~45980 × 45980, 15.8GB). The number of samplings (*K*) is set to 100 and the size is set according to *Max_size*/*K* where *Max_size* = 45980/2. Next, for both *X* and *Y*, we combine the sampled datapoints in a vector (*sample_X*, *sample_Y*) and calculate the correlation between *sample_X* and *sample_Y* by linear regression fitting using Ordinary Least Squares.

We obtained a p-value of the fitted regression lines from t-tests with the null hypothesis that the line is horizontal.

We further evaluated the null hypothesis that slopes and intercepts of the best fitting lines of empirical and null-model data are equal. We performed the ANCOVA test [McDonald, 2009], which evaluates the significance of an F-test under the null hypothesis that both slopes and intercepts are equal. To correct for over-representation of datapoints in these tests, we corrected the degrees of freedom to be (*size_XY* – *C*), where *size_XY* is the number of datapoints and *C* is the number of parameters. For the linear regression fitting, *C* equals to 2 (intercept and slope of best fitting line). For the ANCOVA test, *C* equals to 4 (intercept and slope of one best fitting line and the difference of these between both best-fitting lines).

**XVI. Gene ontology (GO) analysis**

To study the GO representation [Ashburner et al., 2000; Gene Ontology Consortium, 2021] of CSR genes, we performed an overrepresentation test using the PANTHER Classification System [Mi et al., 2019]. This test finds statistically significant overrepresentations using Fisher's exact test, which rejects the null hypothesis that there are no associations between the genes' cohort and the corresponding GO of the biological process for p-values < 0.05. This p-value is corrected for the False Discovery Rate (FDR) using the Benjamini-Hochberg procedure [Benjamini and Hochberg, 1995].

**XVII. Analysis of gene fitness**

From 4133 reference bacterial DNAs with listed genes [Xavier et al., 2021], we used the Rentrez package [Winter et al., 2017] to count in how many of these genomes one finds each gene of MG1655 (GCF_000005845.2_ASM584v2). We use these numbers (divided by the total number of genomes) as a measure of the evolutionary fitness of each gene, which in bacteria can propagate in the biosphere by cell division or by horizontal gene transfer. In detail, we calculated the mean and $CV^2$ of gene fitness of CSR genes as well as for all genes in the genome, along with their standard errors using bootstrapping (104 resampling with replacement). For mixed genes cohort, we calculated the same along with their standard errors using bootstrapping (104 resampling with replacement) but here each sample consists of genes from different groups of sampled at different sizes.

**XVIII. Short-term responses to CS cannot be explained by Transcription Factor interactions**

We studied if the dynamics of CSR genes during CS is influenced by their direct input TFs. According to Regulon DB [Santos-Zavaleta A et al., 2019], the 4698 genes of *E. coli* have a total of 4590 TF interactions between them. Our RNA-seq data informs on the $LFC_{CS}$ of most genes (4328), which have 4435 TF interactions between them. These genes include all 381 CSR genes identified. Also included are 733 TF interactions between them (Supplementary File X3).

Given the time length of protein assembly and maturation [Balleza et al., 2018; Hebisch et. al., 2013; Maurizi, 1992], we expect a time delay from the moment RNAs change until the moment the corresponding TFs change. Once this happens, we expect a short time delay until their output genes' RNA numbers to change in response [Bernstein et al., 2002]. Thereby, we searched for correlations between RNA numbers of genes coding for a TF in a given time moment, and the RNA numbers in a future moment of genes whose promoters are known outputs of that TF (information of the TF-promoter interactions was obtained from Regulon DB).

From the time-lapse RNA-seq data (Supplementary Section I), neither the mid- nor the long-term responses of CSR genes (at 80 and 180 min after CS, respectively) are correlated with the short-term changes (20 min after CS) in the numbers of the RNAs coding for their input TFs.

It is worth noting that, although CSR genes do not show influence from their TFs, there is a visible propagation of information in the TFN during CS. Namely, on average, non-CSR genes exhibited correlated dynamics with their input TFs (Supplementary Figures S17). As such, the lack of correlation during CS is a feature of CSR genes, rather than genome wide. We observed the same at optimal temperatures (Supplementary Figure S17). Potentially, these interactions will be active under conditions not studied here.

Finally, we searched for (but failed to find) correlations between the short-term responses to CS of global transcription regulators (Supplementary Table S5 and Supplementary Figure S1).

**XIX. Short-term responses of CSR genes cannot be explained by the promoters' AT richness**

For CSR genes to exhibit strong repression upon CS, they necessarily were strongly expressing in optimal conditions. This is confirmed in Supplementary Figures S20A (and agrees with data in [Taniguchi et al., 2010]).

Meanwhile, AT-rich promoters are more strongly expressed than GC rich promoters [Liu et al., 2004] in optimal conditions. This could imply that CSR promoters are AT-rich. We thus confronted the AT richness (Supplementary section XIII) of promoter sequences with their short-term response strengths to CS (Supplementary Figures S20B). While we find a genome-wide correlation, that correlation does not exist in the CSR gene cohort (Supplementary Figures S20B inset). We conclude that AT richness is not involved in short-term responsiveness to CS.

### Supplementary Figures

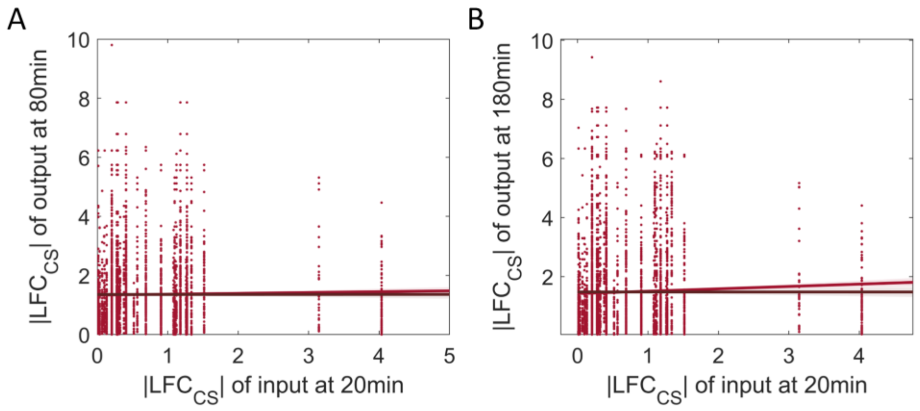


**Figure S1:** Signal propagation after cold shock. Correlation plots of the |LFC$_{CS}$| between input and output genes with a transcription factor pathlength of 1. **(A)** Correlation during CS between output genes at 80min vs input genes at 20min of all pairs of genes input-output (blue dots) and of pairs where the input is a global regulator (dark red dots). **(B)** Correlation during CS between output genes at 180min vs input genes at 20 min of all pairs of genes input-output (blue dots) and of pairs where the input is a global regulator (dark red dots). To the red circles, we fitted by OLS the best fit line. Null models were generated as described in Supplementary Section XV. For each time point, we did an ANOVA test to test for the null hypothesis that the red and black line are not statistically distinguishable. P-values are presented in Supplementary Table S4. For each plot fitted line we performed a likelihood ratio test between the zero-order polynomial and higher order polynomials. P-values < 0.05 reject the null hypothesis that the best fit line is a horizontal line. Shadow areas are 68% confidence bounds.

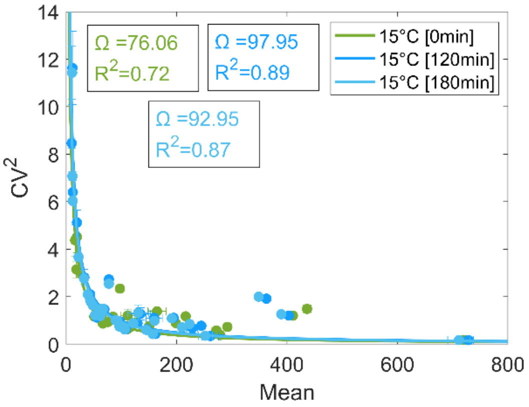


**Figure S2:** Squared Coefficient of variation ($CV^2$) as a function of Mean Protein Fluorescence level. **(A)** $CV^2$ versus mean protein fluorescence level measured by the FITC-H channel of the flow cytometer at 15 °C, 0 min, 120 min and 180 min. To the data we fit the best-fit function $CV^2 = \Omega / P$ [Bar-Even et al., 2006; Taniguchi et al., 2010], where **Ω** is a constant (values for each curve are shown in the insets) and P is the mean protein numbers. We performed a 2-sample t-test to check for the null hypothesis that there is no difference between the Ω values at 120min and 180min. The test does not reject the null hypothesis with a p-value of 0.43.

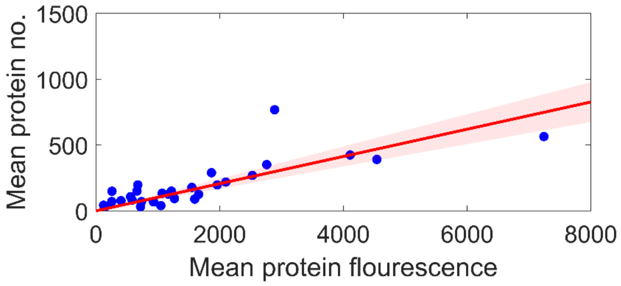


**Figure S3**: Our measured mean protein fluorescences (Supplementary section II) are plotted against the corresponding protein numbers reported in [Taniguchi et al., 2010] in the same growth conditions. The best fitting line of the type y=mx has a statistical significant slope m = 0.10 and its $R^2$=0.56. We performed a F-test on the regression model, which tests for the hypothesis that the 0 order polynomial fits significantly better than a 1st order polynomial. The test rejected the null hypothesis (p-value > $10^{-6}$).

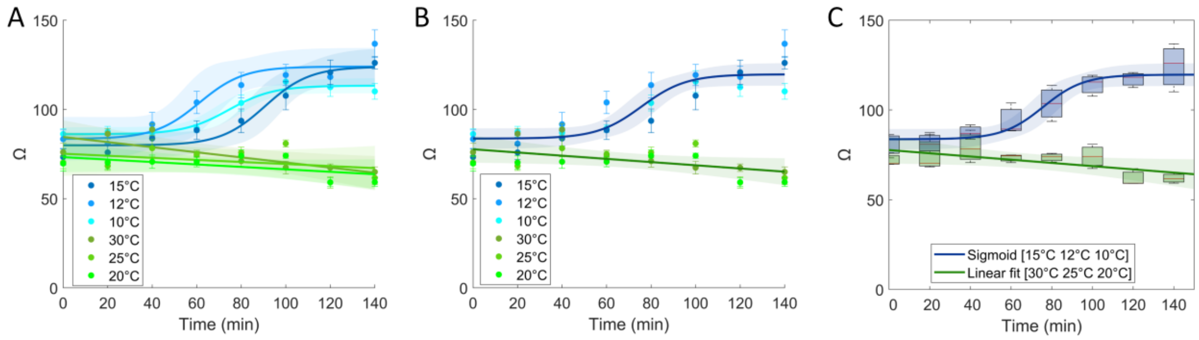


**Figure S4: (A)** For temperature we fit the best fitting function. Data >= 20°C is best fit by a 1st order polynomial while data < 20°C is best fit by a sigmoid function (function given by $\frac{L}{1+e^{-0.1\cdot(x-x_0)}}$, where $L$ is the curve's maximum value and $x_0$ is the $x$ value of the sigmoid midpoint). **(B)** Average best fitting functions for data below 20°C and above or equal to 20°C. The dark blue and green lines correspond to the best fit curves (that maximize $R^2$), for the 15°C-10°C dataset and for the 30°C - 20°C dataset, respectively. **(C)** Box plot of Ω as a function of time for temperature set (Control and CS). The red line in the box is the median and the top and bottom of the box are one STD above and below the median, respectively. For control and cold-shock temperatures, we fit the best fitting function.

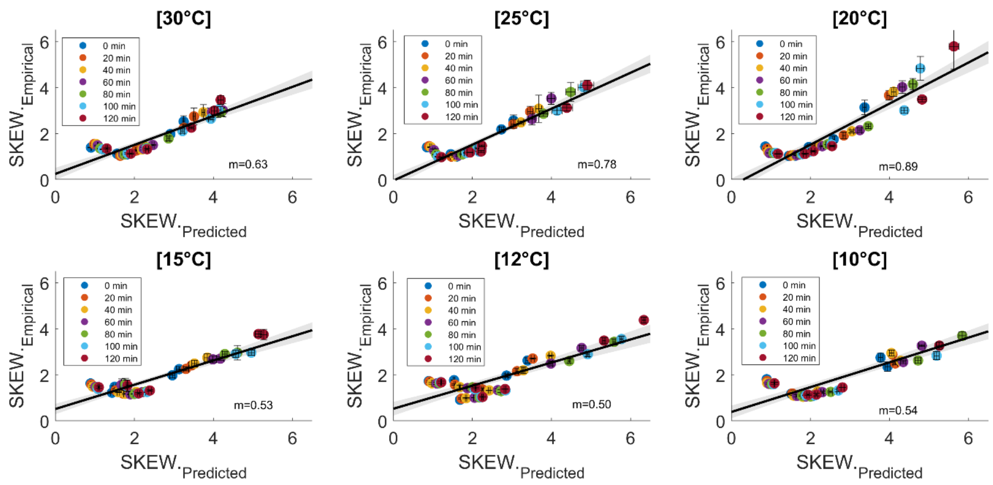


**Figure S5:** Correlation between empirical and predicted skewness for 30°C, 25°C, 20°C, 15°C, 12°C, 10°C. Predicted skewness is estimated from the Ω estimated from the relationship $CV^2$ vs M (Supplementary section XIV) from empirical Flow cytometry data while the empirical skewness is the skewness of the empirical distributions obtained by Flow cytometry. For all fittings $R^2$ is > 0.8.

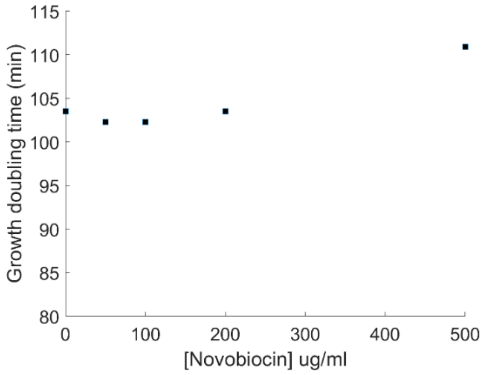


**Figure S6:** *E. coli* K-12 MG1655 cell growth doubling time versus various concentrations of Novobiocin in M9 medium supplemented with 0.4% glucose, amino acids, and vitamins at 30 ºC. The doubling time was measured by the initial $OD_{600nm}$ value, the final $OD_{600\ nm}$ value and the time interval in between.

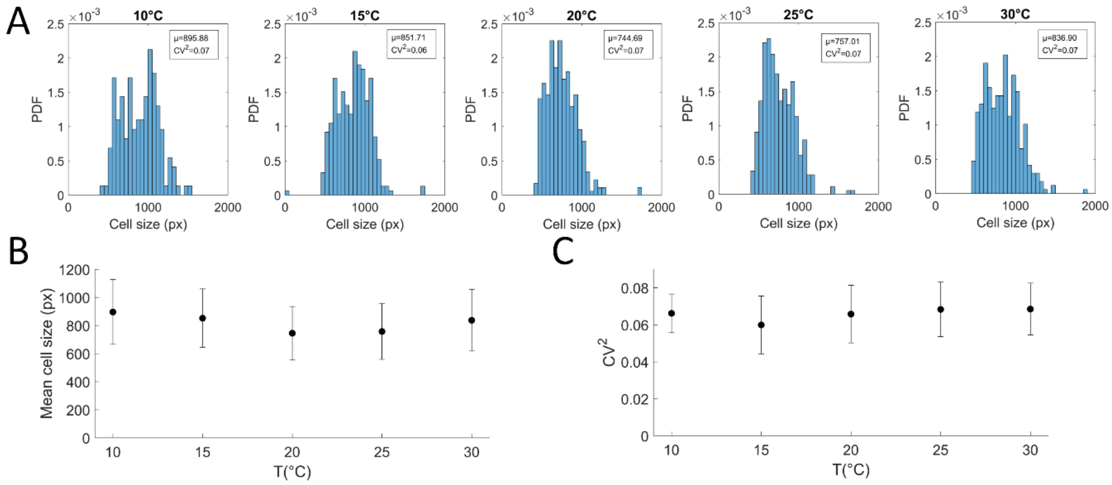


**Figure S7:** Microscopy measurements of cell size (px) of *E. coli* cells expressing the manY gene endogenously tagged with YFP coding sequence at different temperatures (10 °C, 15 °C, 20 °C, 25 °C and 30 °C). **(A)** Probability density function (PDF) of the distribution of cell size, 180 min after the temperature shift. **(B)** Mean cell size (px) as a function of temperature. Vertical error bars correspond to the standard deviation. **(C)** Squared coefficient of variation as a function of temperature. Vertical error bars were calculated by bootstrapping (standard deviation of 500 resamples of 20 cells each).

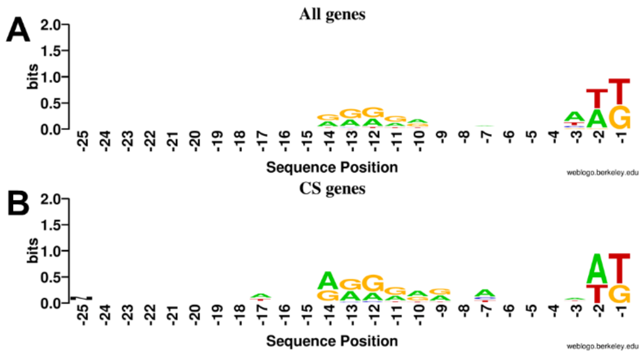


**Figure S8:** Sequence logos of the RBS sequence from the SD to the start codon for **(A)** 179 genes and **(B)** and the 31 CS genes of those 179. Data from RegulonBD. The last nucleotide of the start codon is placed in the position -1. Also, the nucleotides positions decrease in the upstream direction.

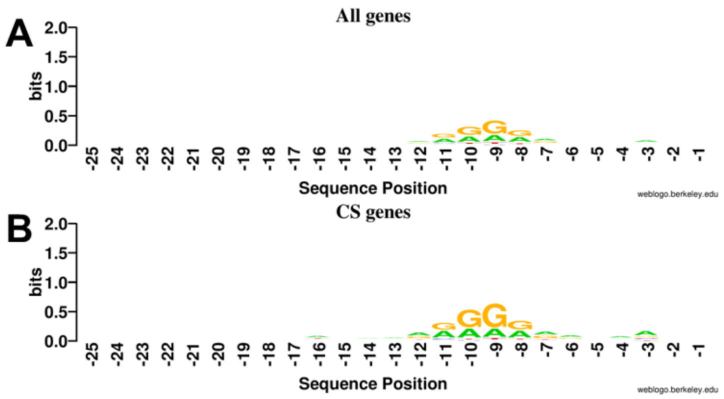


**Figure S9:** Sequence logo of the RBS sequence ranging from the SD sequence to the start codon for all 4357 genes **(A)** and 377 CS genes **(B)** in the other list. The nucleotide upstream of the start codon of the gene is assumed to have the position -1 and the position of a nucleotide decreases as it is located more upstream.

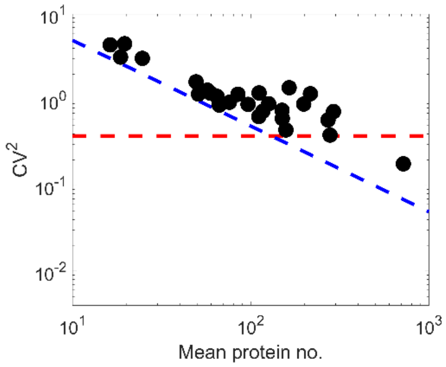


**Figure S10:** Squared coefficient of variation versus mean protein numbers. For mean protein number < 150 the protein expression noise is inversely proportional to the mean with a lower noise limit (blue dashed line), corresponding to intrinsic noise [Taniguchi et al., 2010]. For mean protein number >150 noise becomes independent of the mean with a lower bound of ~0.38 (red dashed line).

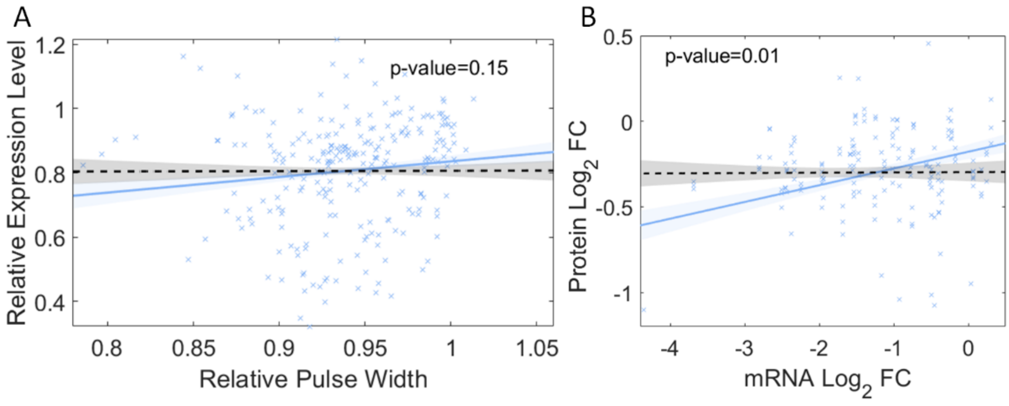


**Figure S11: (A)** Correlation plot between the pulse width and protein expression levels, measured by flow cytometer (Supplementary Section II). Values are relative to 0 min. Black dashed line is the null model (Supplementary Section XV), the two lines are not statistical distinguishable (p-value = 0.15). **(B)** Correlation plot of the |$LFC_{CS}$| of mRNAs (measured by RNAseq at 15 °C, at time 20 min and 80 min after the temperature shift) and protein expression levels (measured by flow cytometry at 15 °C, 120 min and 180 min after (with a gap of 100 min [Startceva et al., 2019]) the temperature shift). We fit the first-order polynomial to the data points by ordinary least-squares and performed an F-test on the regression model, which tests for the hypothesis that the 0 order polynomial fits significantly better than a 1st order polynomial. The test rejected the null hypothesis (p-value = 0.0002). Black dashed line is the null model, the two lines are statistical distinguishable (p-value = 0.01) (Supplementary Section XV). This is consistent with the cell division times (~150 min in M9 media). As such, we expect phenotypic consequences from the cold-shock response at the RNA level to be propagated into the proteins produced in the same period [Newman, et al., 2006; Vogel, et al., 2012; Liu, et al., 2016].

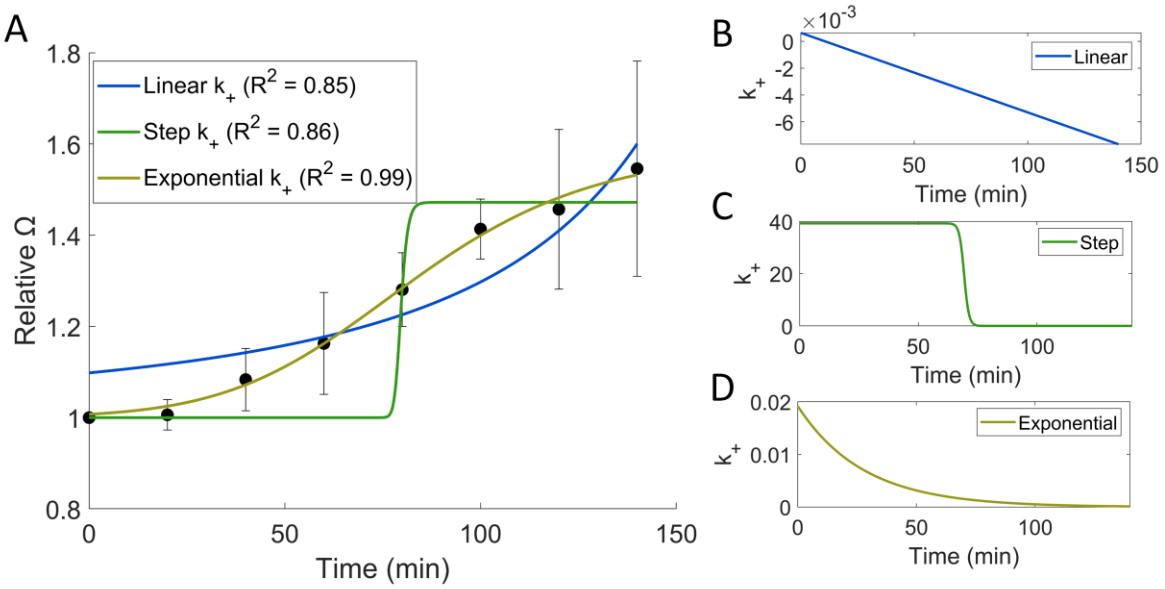


**Figure S12: (A)** Model fitting given a specific function for $k_+$ changes (linear, step and exponential function). Black dots are the empirical data for CS temperatures and error bars are the standard error. **(B-D)** Changes in $k_+$ over time assuming a linear, step and exponential function, respectively. In **A**, **B**, **C**, **D**, all lines are best fitting curves for respective model predictions. Table S12 shows the $R^2$ for each model fit and prediction. Due to low $R^2$ when fitting the empirical data the fitting of quadratic type function is not shown.

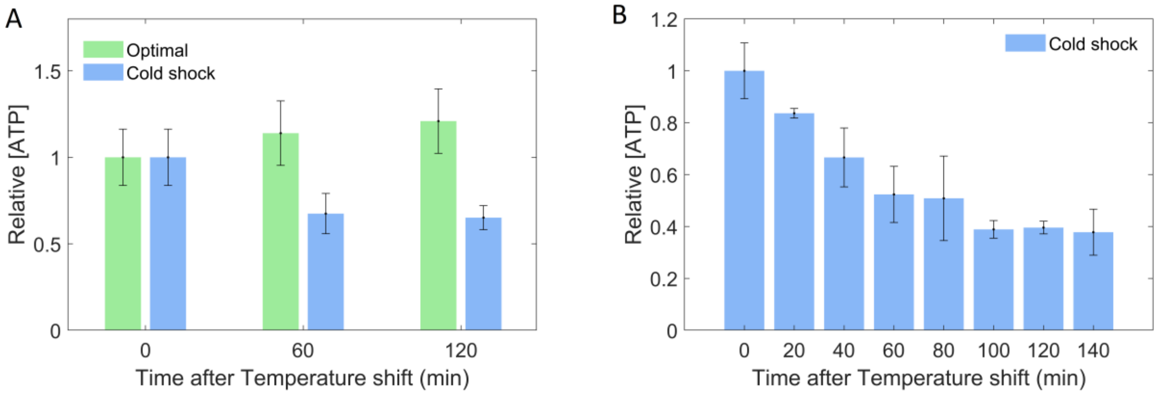


**Figure S13:** Relative ATP concentration measured (Methods Section 2.4) using the QUEEN-2m sensor [Yaginuma et al., 2014], with 3 replicates per condition. Detailed description of correction for autofluorescence and cell division is in (Supplementary Section XII).

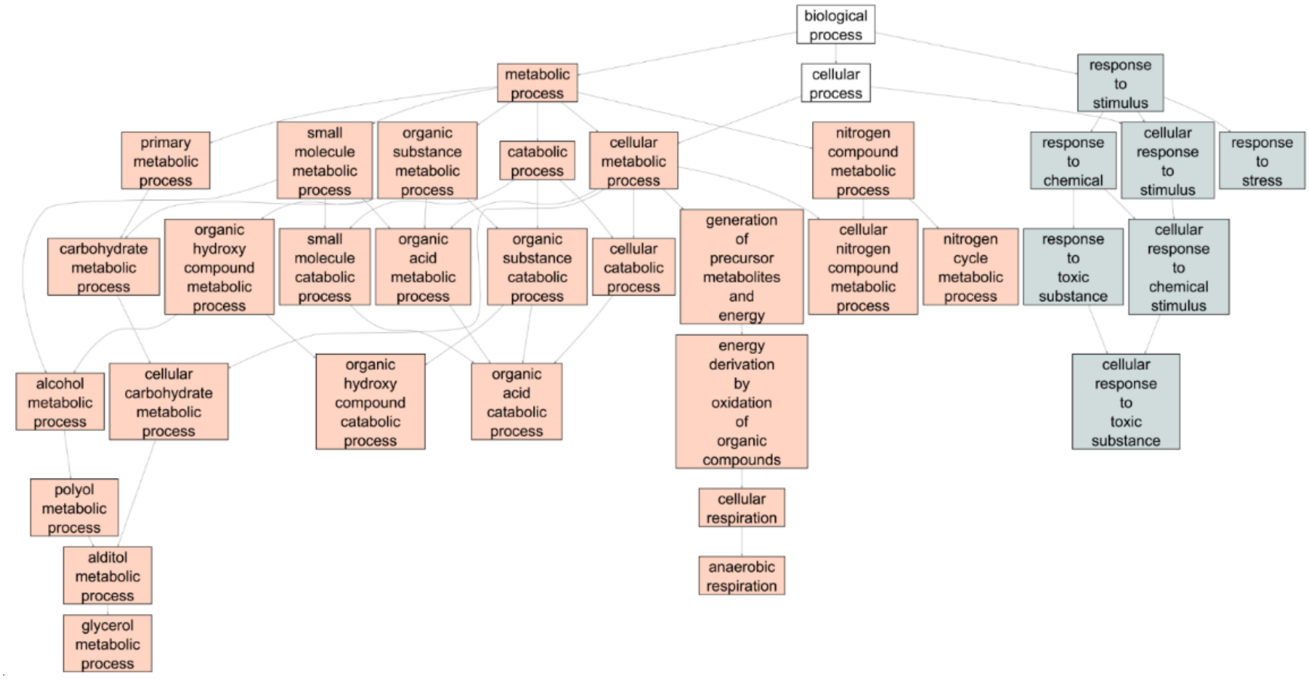


**Figure S14:** Graph of overrepresented gene ontology terms and their ancestors for CSR genes (Supplementary Section XVI). The more general biological processes are connected to specific ontology terms by an arrow pointing to the latter. The ontologies that have the ancestor "metabolic process" and the ones with the ancestor "response to stimulus" are marked separately.

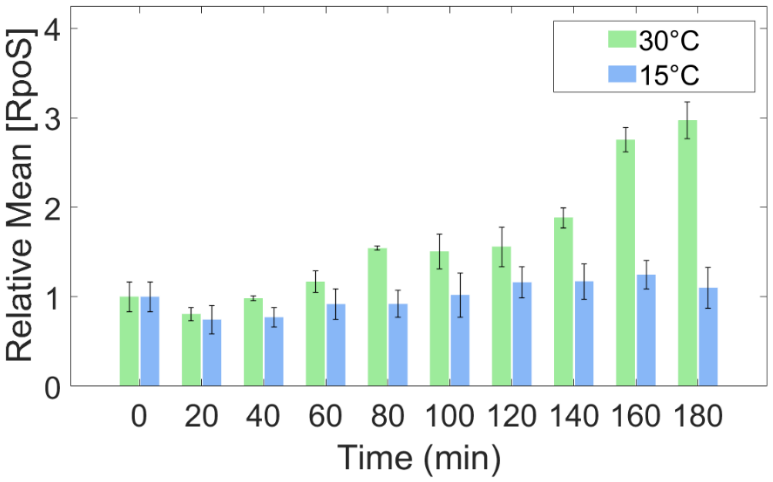


**Figure S15:** Mean concentration of RpoS ($\sigma^{38}$), relative to 0 min, measured every 20 min in both optimal (30 °C) and cold temperature (15 °C), by flow cytometer.

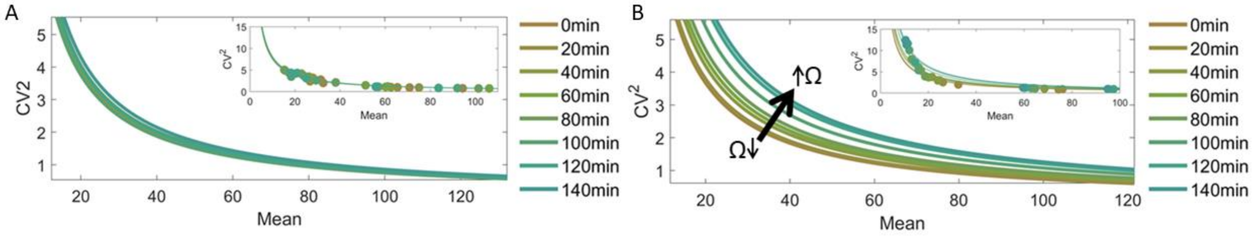


**Figure S16:** $CV^2$ versus mean protein numbers measured by flow cytometer during optimal and CS temperatures, every 20 min for 120 min, respectively.  To each time moment we fit the function $CV^2 = \Omega / M$ [Bar-Even et al., 2006; Taniguchi et al., 2010]. Fittings are shown for each time moment. Each time moment data comes triplicates from 6 genes (aldA, feoA, many, ndk, pepN, tktB). **(A-B inset)**. Data points along with best-fit curves are shown.

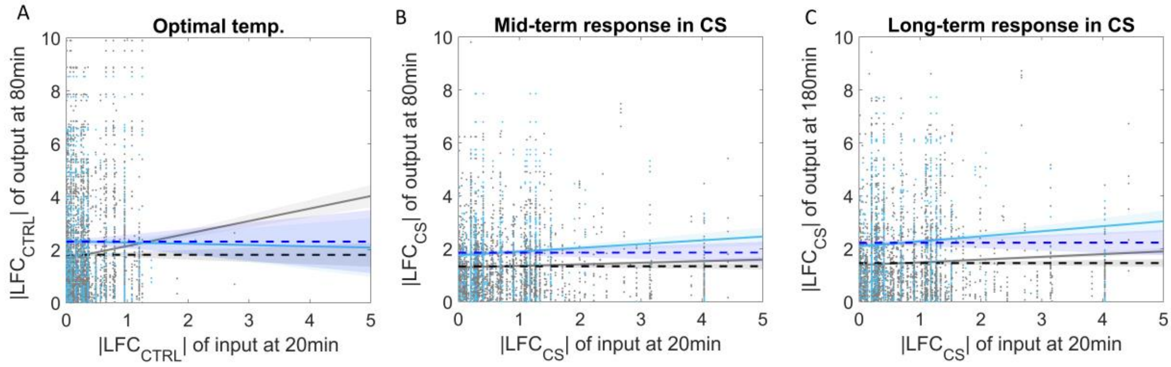

**Figure S17. (A)** Scatter plot between |$LFC_{CS}$| of TF output genes at 80 min and corresponding TF input genes at 20 min in optimal conditions. **(B) and (C)** Scatter plots between |$LFC_{CS}$| of output gene at 80 min and 180 min, respectively, and the corresponding TF input genes at 20 min during CS. Grey circles are all pairs of genes (4435 pairs) (Supplementary section XVIII). Blue circles (733) are of pairs of input-output TF genes for which the output is a CSR gene. We fitted by ordinary least-squares a best-fit line (grey and blue lines, respectively). Dashed lines are the null models (Supplementary Section XV). For each best fit line (grey and blue line) and correspondent null model (black and dark blue dashed line, respectively), we performed an ANCOVA test for the null hypothesis that the two lines are not statistical distinguishable (Supplementary Section XV). P-values < 0.05 reject the null hypothesis. P-values for each test are in Table S2 and Table S3, respectively. Shadow regions are 68% confidence bounds.

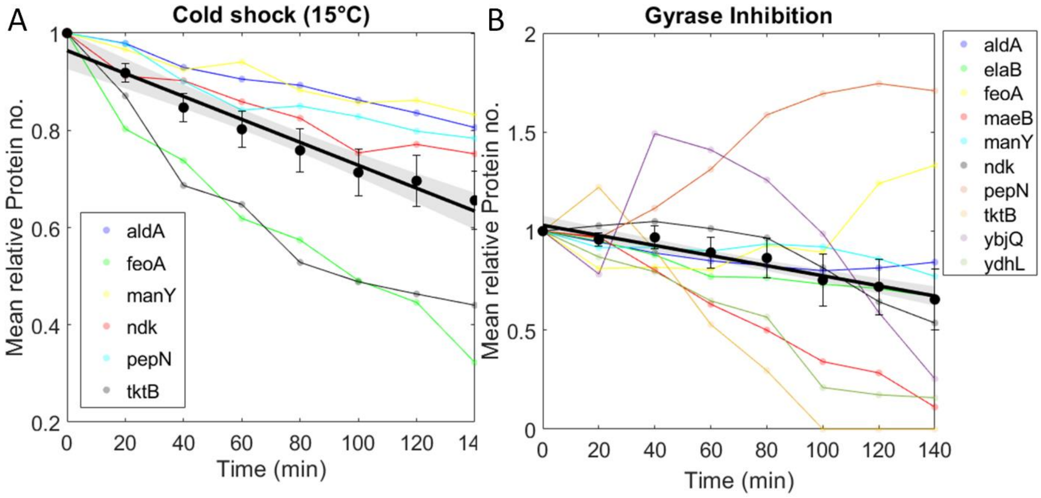


**Figure S18: (A)** Change in relative mean protein numbers of 6 CSR genes over time following CS. **(B)** Change in relative mean protein numbers of 10 CSR genes over time following Gyrase inhibition by Novobiocin. Related with Section 3.7 of the main manuscript.

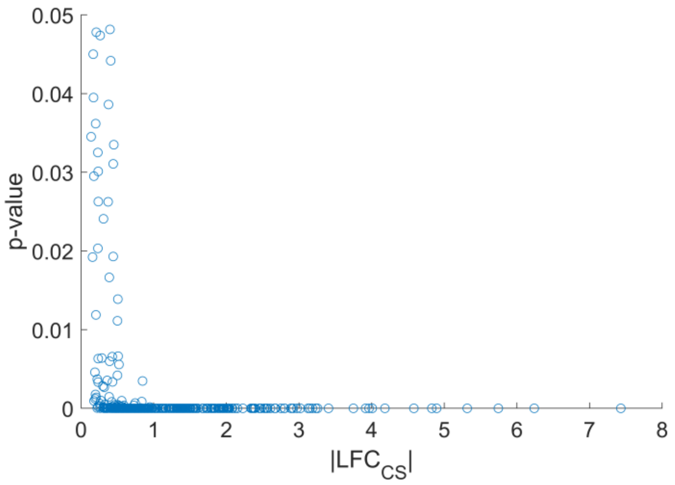


**Figure S19:** Scatter plot between |$LFC_{CS}$| and p-values of the CSR genes from the RNA-seq data.

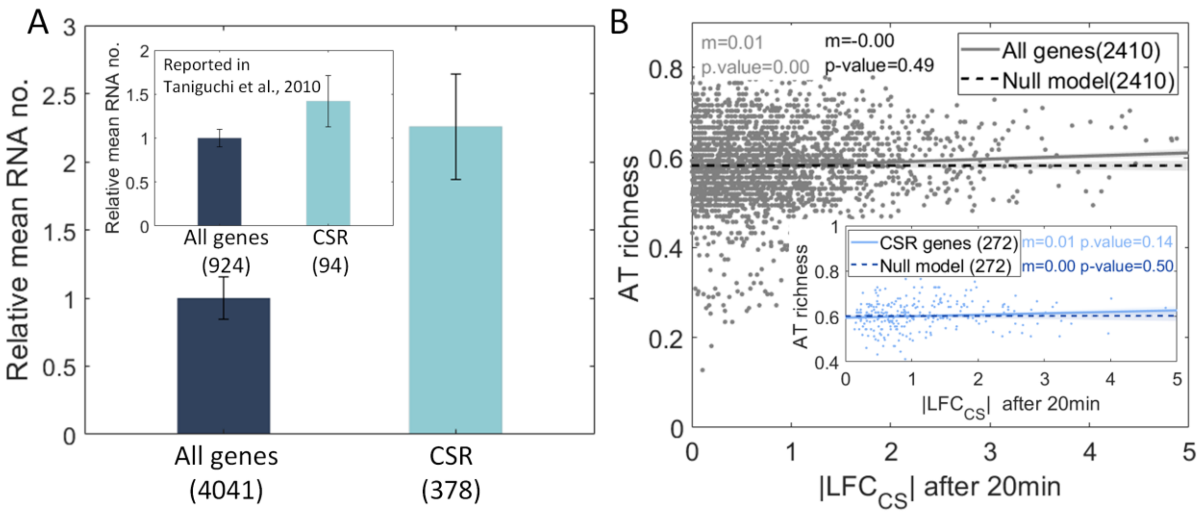


**Figure S20: (A)** Relative RNA expression levels of all genes in the RNAseq (4041) and of the CSR genes alone (378) at optimal temperature. Inset data from [Taniguchi et al., 2010]. **(B)** Correlation between the genes' AT richness and their response to CS (|$LFC_{CS}$|) 20 min after CS. The inset shows the same plot when considering only CSR genes.

**Supplementary Tables**

**Table S1:** List of YFP-fusion strains [Taniguchi et al., 2010] used in this study.

| S. No. | Strain name | Genotype | Source |
|---|---|---|---|
| 1 | MG1655 | λ-, rph-1 | Yale CGSC (CGSC # 6300) |
| 2 | MGmCherry | Same as MG1655, with rpoS::mCherry, chromosomally integrated, replacing rpoS | Gift of James Locke [Patange 2018] |
| 3 | SX1047 | F-, Δ(argF-lac)169, gal-490, Δ(modF-ybhJ)803, λ[cI857 Δ(cro-bioA)], gyrA791-YFP(::cat), IN(rrnD-rrnE)1, rph-1 | Yale CGSC (CGSC # 12602) |
| 4 | SX1056 | F-, Δ(argF-lac)169, gal-490, Δ(modF-ybhJ)803, λ[cI857 Δ(cro-bioA)], IN(rrnD-rrnE)1, rph-1, gyrB792-YFP(::cat) | Yale CGSC (CGSC # 12611) |
| 5 | SX1020 | F-, Δ(argF-lac)169, gal-490, Δ(modF-ybhJ)803, λ[cI857 Δ(cro-bioA)], topA791-YFP(::cat), IN(rrnD-rrnE)1, rph-1 | Yale CGSC (CGSC # 12575) |
| 6 | SX1440 | F-, Δ(argF-lac)169, gal-490, Δ(modF-ybhJ)803, λ[cI857 Δ(cro-bioA)], topB792-YFP(::cat), IN(rrnD-rrnE)1, rph-1 | Yale CGSC (CGSC # 12995) |
| 7 | SX1051 | F-, Δ(argF-lac)169, gal-490, Δ(modF-ybhJ)803, λ[cI857 Δ(cro-bioA)], IN(rrnD-rrnE)1, rph-1, rpoB791-YFP(::cat) | Yale CGSC (CGSC # 12606) |
| 8 | SX1397 | F-, Δ(argF-lac)169, gal-490, Δ(modF-ybhJ)803, λ[cI857 Δ(cro-bioA)], clpA791-YFP(::cat), IN(rrnD-rrnE)1, rph-1 | Yale CGSC (CGSC # 12952) |
| 9 | SX1519 | F-, Δ(argF-lac)169, gal-490, Δ(modF-ybhJ)803, λ[cI857 Δ(cro-bioA)], pepN794-YFP(::cat), IN(rrnD-rrnE)1, rph-1 | Yale CGSC (CGSC # 13074) |
| 10 | SX1812 | F-, yaeH791-YFP(::cat), Δ(argF-lac)169, gal-490, Δ(modF-ybhJ)803, λ[cI857 Δ(cro-bioA)], IN(rrnD-rrnE)1, rph-1 | Yale CGSC (CGSC # 13367) |
| 11 | SX1986 | F-, Δ(argF-lac)169, gal-490, Δ(modF-ybhJ)803, λ[cI857 Δ(cro-bioA)], ydfG791-YFP(::cat), IN(rrnD-rrnE)1, rph-1 | Yale CGSC (CGSC # 13541) |
| 12 | SX1882 | F-, Δ(argF-lac)169, gal-490, Δ(modF-ybhJ)803, λ[cI857 Δ(cro-bioA)], yqjD791-YFP(::cat), IN(rrnD-rrnE)1, rph-1 | Yale CGSC (CGSC # 13437) |
| 13 | SX1394 | F-, Δ(argF-lac)169, gal-490, Δ(modF-ybhJ)803, λ[cI857 Δ(cro-bioA)], ndk-791-YFP(::cat), IN(rrnD-rrnE)1, rph-1 | Yale CGSC (CGSC # 12949) |
| 14 | SX1989 | F-, Δ(argF-lac)169, gal-490, Δ(modF-ybhJ)803, λ[cI857 Δ(cro-bioA)], yeeX793-YFP(::cat), IN(rrnD-rrnE)1, rph-1 | Yale CGSC (CGSC # 13544) |
| 15 | SX1505 | F-, Δ(argF-lac)169, gal-490, Δ(modF-ybhJ)803, λ[cI857 Δ(cro-bioA)], yciI793-YFP(::cat), IN(rrnD-rrnE)1, rph-1 | Yale CGSC (CGSC # 13060) |
| 16 | SX1695 | F-, Δ(argF-lac)169, gal-490, Δ(modF-ybhJ)803, λ[cI857 Δ(cro-bioA)], elaB792-YFP(::cat), IN(rrnD-rrnE)1, rph-1 | Yale CGSC (CGSC # 13250) |
| 17 | SX1950 | F-, Δ(argF-lac)169, gal-490, Δ(modF-ybhJ)803, λ[cI857 | Yale CGSC |

| | | | |
|---|---|---|---|
| | | Δ(cro-bioA)], putP792-YFP(::cat), IN(rrnD-rrnE)1, rph-1 | (CGSC # 13505) |
| 18 | SX1550 | F-, Δ(argF-lac)169, gal-490, Δ(modF-ybhJ)803, λ[cI857 Δ(cro-bioA)], IN(rrnD-rrnE)1, glpD792-YFP(::cat), rph-1 | Yale CGSC (CGSC # 13105) |
| 19 | SX1674 | F-, Δ(argF-lac)169, gal-490, Δ(modF-ybhJ)803, λ[cI857 Δ(cro-bioA)], gcvT792-YFP(::cat), IN(rrnD-rrnE)1, rph-1 | Yale CGSC (CGSC # 13229) |
| 20 | SX1919 | F-, Δ(argF-lac)169, gal-490, Δ(modF-ybhJ)803, λ[cI857 Δ(cro-bioA)], gcvP791-YFP(::cat), IN(rrnD-rrnE)1, rph-1 | Yale CGSC (CGSC # 13474) |
| 21 | SX1917 | F-, Δ(argF-lac)169, gal-490, Δ(modF-ybhJ)803, λ[cI857 Δ(cro-bioA)], gabD791-YFP(::cat), IN(rrnD-rrnE)1, rph-1 | Yale CGSC (CGSC # 13472) |
| 22 | SX1901 | F-, Δ(argF-lac)169, gal-490, Δ(modF-ybhJ)803, λ[cI857 Δ(cro-bioA)], aldA791-YFP(::cat), IN(rrnD-rrnE)1, rph-1 | Yale CGSC (CGSC # 13456) |
| 23 | SX1488 | F-, Δ(argF-lac)169, gal-490, Δ(modF-ybhJ)803, λ[cI857 Δ(cro-bioA)], IN(rrnD-rrnE)1, rph-1, tnaA791-YFP(::cat) | Yale CGSC (CGSC # 13043) |
| 24 | SX1284 | F-, Δ(argF-lac)169, gal-490, Δ(modF-ybhJ)803, λ[cI857 Δ(cro-bioA)], gatZ794-YFP(::cat), IN(rrnD-rrnE)1, rph-1 | Yale CGSC (CGSC # 12839) |
| 25 | SX1763 | F-, Δ(argF-lac)169, gal-490, Δ(modF-ybhJ)803, λ[cI857 Δ(cro-bioA)], manY793-YFP(::cat), IN(rrnD-rrnE)1, rph-1 | Yale CGSC (CGSC # 13318) |
| 26 | SX1416 | F-, Δ(argF-lac)169, gal-490, Δ(modF-ybhJ)803, λ[cI857 Δ(cro-bioA)], pgk-791-YFP(::cat), IN(rrnD-rrnE)1, rph-1 | Yale CGSC (CGSC # 12971) |
| 27 | SX1087 | F-, Δ(argF-lac)169, bolA791-YFP(::cat), gal-490, Δ(modF-ybhJ)803, λ[cI857 Δ(cro-bioA)], IN(rrnD-rrnE)1, rph-1 | Yale CGSC (CGSC # 12642) |
| 28 | SX1526 | F-, Δ(argF-lac)169, gal-490, Δ(modF-ybhJ)803, λ[cI857 Δ(cro-bioA)], katE791-YFP(::cat), IN(rrnD-rrnE)1, rph-1 | Yale CGSC (CGSC # 13081) |
| 29 | SX1771 | F-, Δ(argF-lac)169, gal-490, Δ(modF-ybhJ)803, λ[cI857 Δ(cro-bioA)], nuoE794-YFP(::cat), IN(rrnD-rrnE)1, rph-1 | Yale CGSC (CGSC # 13326) |
| 30 | SX1954 | F-, Δ(argF-lac)169, gal-490, Δ(modF-ybhJ)803, λ[cI857 Δ(cro-bioA)], tktB792-YFP(::cat), IN(rrnD-rrnE)1, rph-1 | Yale CGSC (CGSC # 13509) |
| 31 | SX1859 | F-, Δ(argF-lac)169, gal-490, Δ(modF-ybhJ)803, λ[cI857 Δ(cro-bioA)], IN(rrnD-rrnE)1, rph-1, yjbQ792-YFP(::cat) | Yale CGSC (CGSC # 13414) |
| 32 | SX1781 | F-, Δ(argF-lac)169, gal-490, Δ(modF-ybhJ)803, λ[cI857 Δ(cro-bioA)], IN(rrnD-rrnE)1, feoA791-YFP(::cat), rph-1 | Yale CGSC (CGSC # 13336) |
| 33 | SX1718 | F-, Δ(argF-lac)169, gal-490, Δ(modF-ybhJ)803, λ[cI857 Δ(cro-bioA)], wrbA791-YFP(::cat), IN(rrnD-rrnE)1, rph-1 | Yale CGSC (CGSC # 13273) |
| 34 | SX1975 | F-, Δ(argF-lac)169, gal-490, Δ(modF-ybhJ)803, λ[cI857 Δ(cro-bioA)], yccJ791-YFP(::cat), IN(rrnD-rrnE)1, rph-1 | Yale CGSC (CGSC # 13530) |
| 35 | SX1085 | F-, Δ(argF-lac)169, gal-490, Δ(modF-ybhJ)803, λ[cI857 | Yale CGSC |

| | | | |
|---|---|---|---|
| | | Δ(cro-bioA)], IN(rrnD-rrnE)1, rph-1, tpiA791-YFP(::cat) | (CGSC # 12640) |
| 36 | SX1349 | F-, Δ(argF-lac)169, gal-490, Δ(modF-ybhJ)803, λ[cI857 Δ(cro-bioA)], IN(rrnD-rrnE)1, rph-1, pfkA791-YFP(::cat) | Yale CGSC (CGSC # 12904) |
| 37 | BW25993 | pRSET B_QUEEN-2m_AMPR | Gift from Hiromi Imamura [Yaginuma et al., 2014] |

**Table S2:** For each condition we did an ANCOVA test to test for the null hypothesis that the grey and black line are not statistically distinguishable. P-values are presented below. For conditions that did not reject the null hypothesis (p-values > 0.05).

| CS | | | Optimal |
|---|---|---|---|
| | 80 min | 180 min | 80 min |
| 20 min | 0.028 | 0.0077 | $2.1\times10^{-04}$ |

**Table S3:** For each condition we did an ANCOVA test to test for the null hypothesis that the light blue and dark blue line are not statistically distinguishable. P-values are presented below. For conditions that did not reject the null hypothesis (p-values > 0.05).

| CS | | | Optimal |
|---|---|---|---|
| | 80 min | 180 min | 80 min |
| 20 min | 0.15 | 0.096 | 0.52 |

**Table S4** For each condition we did an ANOVA test (Supplementary Section XV) to test for the null hypothesis that the red and dark red lines are not statistically distinguishable. P-values are presented below. For conditions that did not reject the null hypothesis (p-values > 0.05).

| CS | | |
|---|---|---|
| | 80 min | 180 min |
| 20 min | 0.3625 | 0.1817 |

**Table S5**: List of global regulators (GR) known to regulate 30 or more genes.

| S.No | GR | No. Genes regulated | Genes that code the GR |
|---|---|---|---|
| 1 | CRP | 574 | crp |
| 2 | FNR | 310 | fnr |
| 3 | IHF | 258 | ihfA; ihfB |
| 4 | Fis | 239 | fis |

| | | | |
|---|---|---|---|
| 5 | H-NS | 194 | hns |
| 6 | ArcA | 184 | arcA |
| 7 | NarL | 138 | narL |
| 8 | Fur | 131 | fur |
| 9 | Lrp | 110 | lrp |
| 10 | NsrR | 84 | nsrR |
| 11 | Cra | 82 | cra |
| 12 | FlhDC | 82 | flhD; flhC |
| 13 | CpxR | 71 | cpxR |
| 14 | NarP | 66 | narP |
| 15 | PhoB | 66 | phoB |
| 16 | LexA | 61 | lexA |
| 17 | PhoP | 59 | phoP |
| 18 | NtrC | 56 | glnG |
| 19 | MarA | 46 | marA |
| 20 | ModE | 46 | modE |
| 21 | SoxS | 42 | soxS |
| 22 | PdhR | 41 | pdhR |
| 23 | NagC | 39 | nagC |
| 24 | ArgR | 38 | argR |
| 25 | OxyR | 35 | oxyR |
| 26 | SlyA | 34 | slyA |
| 27 | IscR | 32 | iscR |
| 28 | CysB | 31 | cysB |
| 29 | PurR | 31 | purR |
| 30 | FhlA | 30 | fhlA |

**Table S6**: Equations of Squared Coefficient of Variation ($CV^2$) and skewness (*S*) as a function of Ω and the mean protein numbers (*M*). Also present is the derivation of $CV^2$ and *S* as a function of the model parameters. Step-by-step derivation in shown in Supplementary Section V

| Variable | Function of Ω and *Mean* | 1-step model | 2-step model | ON-OFF model |
|---|---|---|---|---|
| **$CV^2$** | $CV^2 = \Omega \cdot \frac{1}{M}$ | $CV^2 = \frac{1}{\frac{k_1 \cdot k_2}{\lambda_1 \cdot \lambda_2}} \cdot \left(1 + \frac{k_2}{\lambda_1 + \lambda_2}\right)$ | $CV^2 = \frac{1}{\frac{\left(\frac{1}{k1} + \frac{1}{ki}\right)^{-1} \cdot k_2}{\lambda_1 \cdot \lambda_2}} \cdot \left(1 + \frac{k_2}{\lambda_1 + \lambda_2}\right)$ | $CV^2 = \frac{1}{\frac{k_+}{k_+ + k_-} \cdot \frac{k_1 \cdot k_2}{y_1 \cdot y_2}} \cdot \left(1 + \frac{k_2}{y_1 + y_2}\left(1 + (1-P)\frac{k_1(k_0 + y_1 + y_2)}{(k_0 + y_1)(k_0 + y_2)}\right)\right)$ |
| **S** | $S = \frac{2}{\sqrt{M}} \cdot \sqrt{\Omega}$ | $S = \frac{2 \cdot \sqrt{\left(1 + \frac{k_2}{\lambda_1 + \lambda_2}\right)}}{\sqrt{\frac{k_1 \cdot k_2}{\lambda_1 \cdot \lambda_2}}}$ | $S = \frac{2 \cdot \sqrt{\left(1 + \frac{k_2}{\lambda_1 + \lambda_2}\right)}}{\sqrt{\frac{\left(\frac{1}{k1} + \frac{1}{ki}\right)^{-1} \cdot k_2}{\lambda_1 \cdot \lambda_2}}}$ | $S = \frac{2 \cdot \sqrt{\left(1 + \frac{k_2}{y_1 + y_2}\left(1 + (1-P)\frac{k_1(k_0 + y_1 + y_2)}{(k_0 + y_1)(k_0 + y_2)}\right)\right)}}{\sqrt{\frac{k_+}{k_+ + k_-} \cdot \frac{k_1 \cdot k_2}{y_1 \cdot y_2}}}$ |

**Table S7:** Reference parameter values assuming the one-step and the ON-OFF models. Given that Lac is considered to be a strong promoter, combining these parameter values should result in relatively high RNA and protein numbers.

| Rate Constant | Description | Values ($s^{-1}$) | References |
|---|---|---|---|
| $k_+$ | Promoter unlocking | $7\times10^{-4}$ | For the native Lac promoter [Chong et al., 2014; Palma et al., 2020] |
| $k_-$ | Promoter locking | 0.0012 | For the native Lac promoter [Stracy et al., 2019; Palma et al., 2020] |
| $k_1$ | Transcription rate | $8.3\times10^{-4}$ to 0.02 | Average expressing native genes [Prajapat et al., 2018] |
| $k_2$ | Translation rate | 0.04 to 0.06 | Average expressing native genes [Bremer et al., 1996] |
| $\lambda_1$ | RNA degradation | 0.004 | Average RNA degradation rates [Bernstein et al., 2002] |
| $\lambda_2$ | Protein decay (degradation and dilution in cell division) | $2.93\times10^{-5}$ | Average protein degradation rates [Koch and Levy 1955; Taniguchi et al., 2010] |

**Table S8:** Set of parameter values use to investigate the noise levels of each model: 1-step model (Model 1 in Figure 8), 2-step model (Model in figure S8) and ON-OFF model (Model 2 in Figure 8)

| Rate Constant | One rate-limiting step model | Two rate-limiting step model | ON-OFF model |
|---|---|---|---|
| $k_1$ ($s^{-1}$) | 0.002 | 0.004 | 0.0057 |
| $k_+$($s^{-1}$) | | | 0.0018 |
| $k_-$($s^{-1}$) | | | 0.0033 |
| $k_i$($s^{-1}$) | | 0.004 | |
| $k_2$($s^{-1}$) | 0.23 | 0.23 | 0.23 |
| $\lambda_1$($s^{-1}$) | 0.004 | 0.004 | 0.004 |
| $\lambda_2$($s^{-1}$) | $2.9\times10^{-5}$ | $2.9\times10^{-5}$ | $2.9\times10^{-5}$ |

**Table S9:** p-distances to the consensus SD sequence (AGGAGG) [Saito et al., 2020]. The sequences were extracted from the available ones in RegulonDB. Shown are the mean and standard deviation of the p-distances of all genes (179 genes), of the 31 genes of those 179 that were found to CSR, and of 1000 cohorts of 31 genes assembled by random selection from the 179 genes.

| | Consensus SD sequence ('AGGAGG') (Mean ± STD) |
|---|---|
| Genome wide p-distance (179 genes) ± STD | 0.57 ± 0.22 |
| p-distance of CSR genes (31 genes) ± STD | 0.48 ± 0.20 |
| p-distance of randomly selected genes (31 genes) | 0.54 ± 0.20 |

**Table S10:** p-distances of the start codon sequences of CSR genes to the most frequent start codon sequences (ATG; GTG; TTG; ATT; CTG) [Blattner et al., 1977; Sacerdot et al., 1982; Missiakas et al.,1993]. Sequences extracted from *Regulon DB.* Shown are the mean and standard deviation of the p-distances of all genes (179 genes), of the 31 genes of those 179 that were found to CSR, and of 1000 cohorts of 31 genes assembled by random selection from the 179 genes.

| | ATG | GTG | TTG | ATT | CTG |
|---|---|---|---|---|---|
| Average start codon sequence (N=179) ± STD *(Regulon DB)* | 0.49 ± 0.22 | 0.64 ± 0.34 | 0.61 ± 0.29 | 0.47 ± 0.18 | 0.63 ± 0.31 |
| Average start codon sequence of CS genes (N=31) ± STD | 0.59 ± 0.47 | 0.71 ± 0.34 | 0.66 ± 0.29 | 0.52 ± 0.17 | 0.67 ± 0.30 |
| Average start codon sequence of random cohort (31) ± STD | 0.53 ± 0.49 | 0.63 ± 0.35 | 0.59 ± 0.28 | 0.52 ± 0.17 | 0.63 ± 0.31 |

**Table S11:** Over-represented biological process according to the Gene Ontology (GO) Overrepresentation Test (Supplementary Section XVI) with Fisher's exact test with FDR correction. For each GO biological process it is shown the number of genes related to it, in the *E. coli* genome (out of the 4390 genes); the number of genes related in the CSR cohort (out of the 376 recognized genes by GO); the number of genes expected to be present in a cohort of the size of the CSR cohort; and finally, the fold-enrichment.

| **Biological Processes** | **GO biological process complete** | **No. Genes in E. coli genome** | **No. Genes in CS cohort** | **Expected No. Genes in CS cohort** | **Fold Enrichment** |
|---|---|---|---|---|---|
| **Metabolic process (GO: 0008152)** | Glycerol metabolic process (GO:0006071) | 20 | 12 | 1,71 | 7,01 |
| | Nitrogen cycle metabolic process (GO:0071941) | 25 | 13 | 2,14 | 6,07 |
| | Organic hydroxy compound catabolic process (GO:1901616) | 52 | 16 | 4,45 | 3,59 |
| | Generation of precursor metabolites and energy (GO:0006091) | 210 | 60 | 17,99 | 3,34 |
| | Organic acid catabolic process (GO:0016054) | 191 | 35 | 16,36 | 2,14 |
| | Carbohydrate metabolic process (GO:0005975) | 389 | 61 | 33,32 | 1,83 |
| | Organic acid metabolic process (GO:0006082) | 542 | 77 | 46,42 | 1,66 |
| **Response to stimulus** | Response to stress (GO:0006950) | 556 | 74 | 47,62 | 1,55 |

| | | | | | |
|---|---|---|---|---|---|
| **(GO: 0050896)** | Cellular response to toxic substance (GO:0097237) | 28 | 10 | 2,4 | 4,17 |

**Table S12:** Best fitting parameter values for model functions of $k_+$ as a function of temperature.

| Function type | Equation | Estimated parameters | Average $R^2$ |
|---|---|---|---|
| Linear | $b_2 \cdot x + b_1$ | $b_1 = 6\times10^{-4}$; $b_2 = -5.9\times10^{-5}$; $k_- = 0.013\ s^{-1}$ | 0.85 |
| Exponential | $e^{(-b\cdot x)} \cdot a$ | b = 0.036; a = 0.019; $k_- = 0.002\ s^{-1}$ | 0.99 |
| Quadratic | $b_3 \cdot x^2 + b_2 \cdot x + b_1$ | $b_1 = 0.847$; $b_2 = 0.894$; $b_3 = -0.024$; $k_- = 1.183\ s^{-1}$ | < 0 |
| Step | $\left(1-\frac{1}{1+e^{-(x-L)}}\right)\times a$ | a = 39.25; L = 69.32; $k_- = 0.003\ s^{-1}$ | 0.86 |
